# Degron-tagged reporters probe membrane topology and enable the specific labelling of membrane-wrapped structures

**DOI:** 10.1101/442103

**Authors:** Katharina B Beer, Gholamreza Fazeli, Kristyna Judasova, Linda Irmisch, Jona Causemann, Jörg Mansfeld, Ann M Wehman

**Affiliations:** Rudolf Virchow Center, Julius-Maximilians-Universität Würzburg, Würzburg, Germany; Dresden International Graduate School for Biomedicine and Bioengineering (DIGS-BB), Dresden, Germany; Cell Cycle, Biotechnology Center, Technische Universität Dresden, Dresden, Germany

## Abstract

Visualization of specific organelles in tissues over background fluorescence can be challenging, especially when reporters localize to multiple structures. Instead of trying to identify proteins enriched in specific membrane-wrapped structures, we used a selective degradation approach to remove reporters from the cytoplasm or nucleus of *C. elegans* embryos and mammalian cells. We demonstrate specific labelling of organelles using degron-tagged reporters, including extracellular vesicles, as well as individual neighbouring membranes. These degron-tagged reporters facilitate long-term tracking of released cell debris and cell corpses, even during uptake and phagolysosomal degradation. We further show that degron protection assays can probe the topology of the nuclear envelope and plasma membrane during cell division, giving insight into protein and organelle dynamics. As endogenous and heterologous degrons are used in bacteria, yeast, plants, and animals, degron approaches can enable the specific labelling and tracking of proteins, vesicles, organelles, cell fragments, and cells in many model systems.

## Introduction

Membranes form barriers that separate cells from their environment and separate diverse subcellular compartments so that they can carry out distinct functions within cells^1^. Membranes are also highly dynamic, undergoing fusion and fission events during endocytosis, ectocytosis, and cytokinesis^2,3^. As membrane bilayers are less than 10 nm in diameter, light microscopic techniques struggle to distinguish neighbouring membranes or to tell when membranes have fused to have a closed topology^4^. Therefore, techniques that allow specific labelling of distinct structures are required to visualize membrane dynamics and probe membrane topology in living cells.

Fluorescent reporters binding specific phosphatidylinositol species are popular for studying membrane dynamics^5^, but these lipids are often present on multiple structures, making it hard to distinguish individual organelles. One example is during phagocytosis, when cellular debris or cell corpses are engulfed by the plasma membrane^6^. Both the corpse and engulfing cell plasma membranes contain the same phosphatidylinositol species, making it challenging to distinguish these membranes in living cells. Electron microscopy and super-resolution light microscopy can visualize the few tens of nm that separate the phagosome membrane from the corpse membrane^7^, but these techniques rely on fixation, which makes it challenging to study dynamics. Another example is extracellular vesicles released from cells. The content of vesicles that bud from the plasma membrane by ectocytosis is similar to the membrane and associated cytoplasm that they originate from^8^, which makes it hard to distinguish released vesicles from the plasma membrane of neighbouring cells. The lack of specific markers for extracellular vesicles using conventional reporters has limited our understanding of their cell biology^9^. Thus, new approaches are needed to label specific membranes.

In order to develop reporters that visualize specific membrane structures, we repurposed the cell’s endogenous machinery for selective degradation. We were inspired by protease protection assays using exogenous proteases^10^, but wanted to establish an *in vivo* system that did not require detergent-mediated permeabilization of the plasma membrane. Degrons are degradation motifs that target specific proteins for ubiquitination and degradation, which has led to degron-tagging being used as an alternative loss-of-function approach to RNA interference or genetic knockouts^11^. Degrons recruit ubiquitin ligases to polyubiquitinate target proteins, resulting in the proteasomal degradation of cytosolic targets or the lysosomal degradation of transmembrane targets^12^. Rather than using degron tags for a loss-of-function technique, we used the degradation of degron-tagged reporters in the cytosol to specifically label certain cells, cell fragments, organelles, and vesicles.

To test endogenous degrons, we primarily used the zinc finger 1 (ZF1) degron from the PIE-1 protein to degrade fluorescent reporters in developing *C. elegans* embryos. The ZF1 degron is a 36 amino acid motif recognized by the SOCS-box protein ZIF-1, which binds to the elongin C subunit of an ECS ubiquitin ligase complex^13^. ZIF-1 is expressed in sequential sets of differentiating somatic cells^14^, resulting in a stereotyped pattern of degradation in developing embryos^13^ (Fig. 1a). Fusing the ZF1 degron to a target protein results in degradation within 30 to 45 min of ZIF-1 expression in both embryonic and adult tissues^15^. As an alternate approach, we used the C-terminal phosphodegrons (CTPD) from the *C. elegans* OMA-1 protein^16^. Two threonines in the C-terminus of OMA-1 are phosphorylated after fertilization, leading to recognition of OMA-1 by multiple SCF ubiquitin ligase complexes and rapid proteasomal degradation in embryos at the end of the 1-cell stage^16,17^ (Fig. 1b). We also used a heterologous degron in mammalian cells, the auxin-inducible degron (AID) from plants^18^. The 68 amino acid AID motif from IAA17 is recognized by the F-Box protein TIR1 in the presence of the auxin family of plant hormones^19^. Auxins are cell permeable and TIR1 is able to become part of endogenous SCF ubiquitin ligase complexes in model systems from yeast to mammals in order to ubiquitinate AID-tagged proteins^18^. The AID system is thus a three-component system, allowing temporal control of degradation by addition of auxin hormone and spatial control of degradation by the expression of TIR1 in different cells (Fig. 1c).

**Fig. 1:**
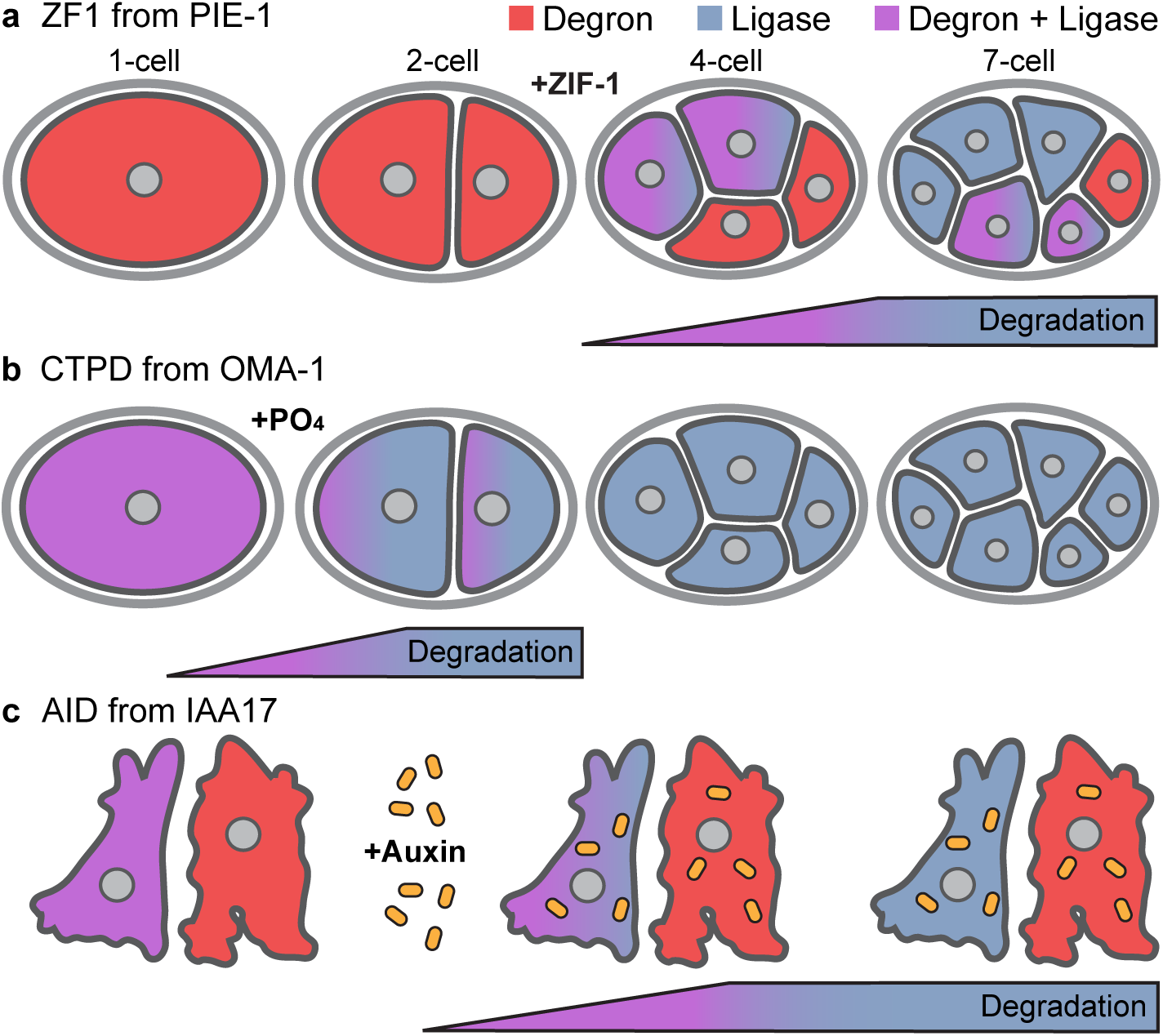
Comparison of degron approaches. a) Proteins with a ZF1 degron (red) are stable before expression of the ubiquitin ligase adaptor ZIF-1 (blue) in *C. elegans* embryos. ZIF starts to be expressed in a stereotyped pattern of somatic cells after the 2-cell stage, starting with the two anterior cells (red + blue = purple). Proteins with a ZF1 tag are degraded in somatic cells, starting with the anterior cells (purple to blue). Cells that do not express ZIF-1, such as the posterior germ cell, do not degrade proteins with a ZF1 tag (red). b) The C-terminal phosphodegrons (CTPD) of OMA-1 are inert until phosphorylated and CTPD-tagged proteins (red) are stable, despite the presence of SCF ubiquitin ligases (blue). Phosphorylation of CTPD occurs during the first mitosis in *C. elegans* embryos, leading to degradation of CTPD-tagged proteins during the first cell division. c) Expression of the TIR1 ligase adaptor (blue) is not sufficient to induce robust degradation of AID-tagged proteins (red). Addition of auxin family hormones induces degradation of AID-tagged proteins in cells where TIR1 is expressed (purple to blue).

Here, we show that degron-tagged reporters separated from the ubiquitin ligase complex by intervening membranes are no longer accessible to ubiquitination and degradation in *C. elegans* embryos or mammalian cells. This results in background-free labelling of specific cells, organelles, and vesicles. This improvement in the signal-to-noise ratio enabled the visualization of extracellular vesicles *in vivo*, the long-term tracking of individual phagosomes, as well as distinguishing a corpse plasma membrane from the engulfing phagosome membrane *in vivo*. In addition, degron-tagging allowed us to measure the timing of nuclear envelope breakdown and abscission during cell division. Degron-tagged reporters thus provide a convenient method for investigating *in vivo* dynamics from the level of proteins to cells.

## Results

To determine whether degron-tagged reporters would be useful for cell biological approaches, we tested whether an endogenous degradation system was capable of degrading abundant reporter proteins and examined whether the increased proteasomal load had negative effects on cells. We tagged a membrane-binding domain, the PH domain of rat PLC1∂1, with the ZF1 degron from *C. elegans* PIE-1 and expressed it in worm embryos. Similar to an mCherry-tagged PH reporter (Fig. 2a), the cytosolic mCh::PH::ZF1 reporter initially localizes to the plasma membrane (Fig. 2d). Thus, the degron tag does not disrupt the normal localization of the reporter.

**Fig. 2:**
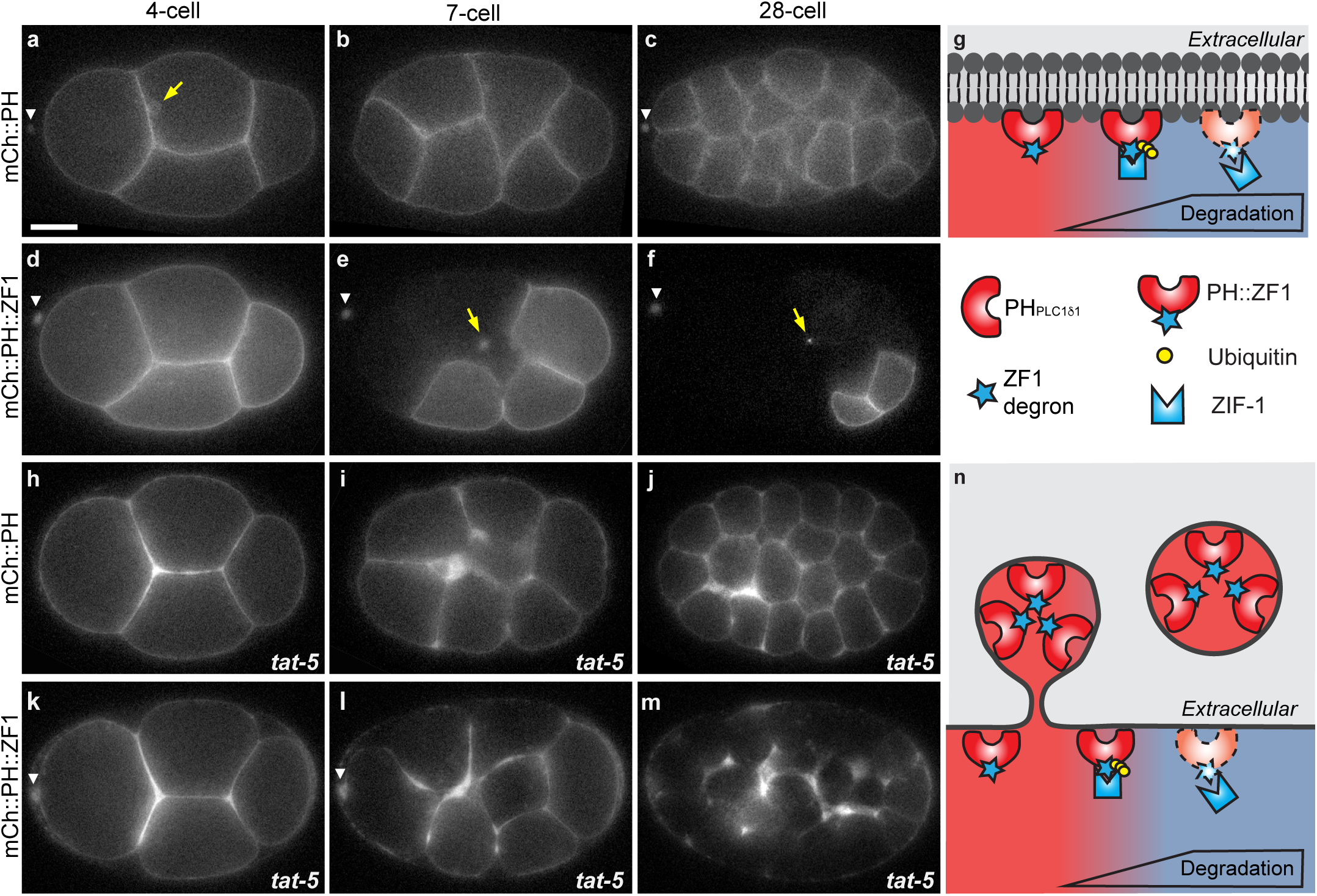
The ZF1 degron enables labelling of specific cells and vesicles in *C. elegans* embryos. a-c) The lipid-binding mCh::PH_PLC1∂1_ reporter localizes to the plasma membrane and endocytic vesicles (arrow) in 4-, 7-, and 28-cell embryos (n=19). Scale bar: 10 µm. d-f) ZIF-1-driven proteasomal degradation of the degron-tagged mCh::PH_PLC1∂1_::ZF1 reporter starts in anterior blastomere (AB) cells during the 4-cell stage (d), leading to the absence of the mCh::PH::ZF1 fluorescence in anterior AB cells at the 7-cell stage (e) and most somatic cells at the 28-cell stage (f, n=10). The ubiquitin ligase adaptor ZIF-1 is not expressed in the posterior germ line or in anterior polar bodies (arrowhead), resulting in the persistence of mCh::PH::ZF1 in these cells. Arrows indicate labelled intracellular vesicles, which are protected from proteasomal degradation by intervening membranes. See also video 1. Anterior is left, dorsal is up. g) The lipid-binding PH domain tagged with the ZF1 degron is recognized by the ubiquitin ligase adaptor ZIF-1 and ubiquitinated. Polyubiquitination leads to proteasomal degradation of degron-tagged fluorescent reporters (dotted lines). h-j) Embryos treated with *tat-5* RNAi show increased mCh::PH membrane labelling due to accumulated microvesicles at the 4-, 7-, and 28-cell stage (n=11). k) Increased mCh::PH::ZF1 is visible at the 4-cell stage in embryos treated with *tat-5* RNAi. l-m) Gradual degradation of mCh::PH::ZF1 in somatic cells facilitates visualization of released microvesicles in a 7- and 28-cell embryo (n=22). Due to degradation of cytosolic mCh::PH::ZF1 reporter at the plasma membrane, even small amounts of released microvesicles are easily visible. See also video 1. n) Degron-tagged PH domain reporters are released in extracellular vesicles that bud from the plasma membrane before ZIF-1 expression. ZIF-1 expression leads to the ubiquitination and proteasomal degradation of cytosolic ZF1-tagged reporters (dotted lines). ZF1-tagged membrane reporters in released vesicles are not ubiquitinated or degraded and maintain fluorescence.

ZF1-mediated degradation begins in anterior somatic cells at the 4-cell stage, due to the onset of expression of the ubiquitin ligase adaptor protein ZIF-1^14^. While mCh::PH fluorescence persists in developing embryos (Fig. 2b-c), mCh::PH::ZF1 is progressively degraded, starting with anterior somatic cells (Fig. 2e-g, video 1). ZIF-1 is not expressed in the germ lineage, resulting in persistent fluorescence in a couple of posterior cells (Fig. 2e-f). ZIF-1 expression also does not occur in two small cell corpses born at the anterior side of the embryo during meiosis^14,20^. These polar bodies maintain mCh::PH::ZF1 fluorescence (arrowheads in Fig. 2). Thus, the degron tag led to rapid degradation of a highly-expressed, exogenous reporter in cells where the ligase adaptor was expressed and the reporter could therefore be ubiquitinated.

As ZIF-1 has a number of known targets whose proteasomal degradation is important for embryonic development^13^, we tested whether the expression of ZF1-tagged reporters disrupted development. Stable transgenic strains expressing various ZF1-tagged reporters were fertile and had similar numbers of viable progeny that did not show developmental delays (Fig. S1a). In fact, most ZF1-tagged reporter embryos developed significantly faster than corresponding reporter strains without the degron (Fig. S1b) and were less likely to show overexpression defects (Fig. S1c-f). This suggests that degron reporters can be tolerated better than other overexpressed reporters and do not generally disrupt development.

### Degron reporters specifically label released extracellular vesicles

In addition to labelling the plasma membrane, mCh::PH and mCh::PH::ZF1 labelled intracellular vesicles (arrows in Fig. 2), some of which maintained their fluorescence in the mCh::PH::ZF1 strain (video 1). As mCh::PH::ZF1 on the cytosolic face of vesicles would be accessible for ubiquitination and proteasomal degradation, the persistence of the degron reporter suggests that it is protected from degradation by intervening membranes. We hypothesized that the PH::ZF1 reporter persisted in extracellular vesicles (EV) or other cell debris that are taken up by the cell during endocytosis.

To test whether the degron-tagged PH reporter could be used to specifically label and track EVs released *in vivo*, we examined microvesicles. Microvesicles are 90-500 nm vesicles that arise from plasma membrane budding, aka ectocytosis^8^. In wild type embryos, microvesicles are difficult to detect due to their low abundance and proximity to the plasma membrane. Microvesicle budding is normally inhibited by the TAT-5 lipid flippase, resulting in continuous microvesicle release when *tat-5* is knocked down, as demonstrated by electron tomography^21^. In mCh::PH embryos, microvesicle overproduction is visible as thickened membrane labelling between *tat-5* knockdown cells (Fig. 2h-j) in comparison to control embryos (Fig. 2a-c). However, small patches of microvesicles are difficult to detect over the background of the plasma membrane fluorescence. In contrast, released microvesicles are clearly visible after *tat-5* knockdown using the mCh::PH::ZF1 reporter (Fig. 2l-m)^2^, due to proteasomal degradation of the plasma membrane label (Fig. 2n). Released EVs are then visible floating between the embryo and the eggshell (Video 1). Thus, degron tagging a general plasma membrane reporter reveals microvesicles and their movement *in vivo*.

To determine whether this was a specific feature of the ZF1 degron or the ECS ubiquitin ligase, we tested whether another degron could be used with SCF ubiquitin ligases to label EVs. We tagged the PH reporter with a C-terminal fragment of OMA-1 (aa219-378) containing two phosphorylation sites important for degradation^16^, which we named the C-terminal phosphodegrons (CTPD) (Fig. 3d). Early during the 1-cell stage, mCh::PH::CTPD localized brightly to the plasma membrane, but began to be degraded during the first mitotic division (Fig. 3a, Video 2). Degradation was transient and some mCh::PH::CTPD persisted on the plasma membrane after the 2-cell stage (Fig. 3c). Even with partial degradation, microvesicles could readily be observed with mCh::PH::CTPD after *tat-5* RNAi treatment (Fig. 3b, Video 2). Thus, even a partial loss of plasma membrane signal enhanced visualization of EVs.

**Fig. 3:**
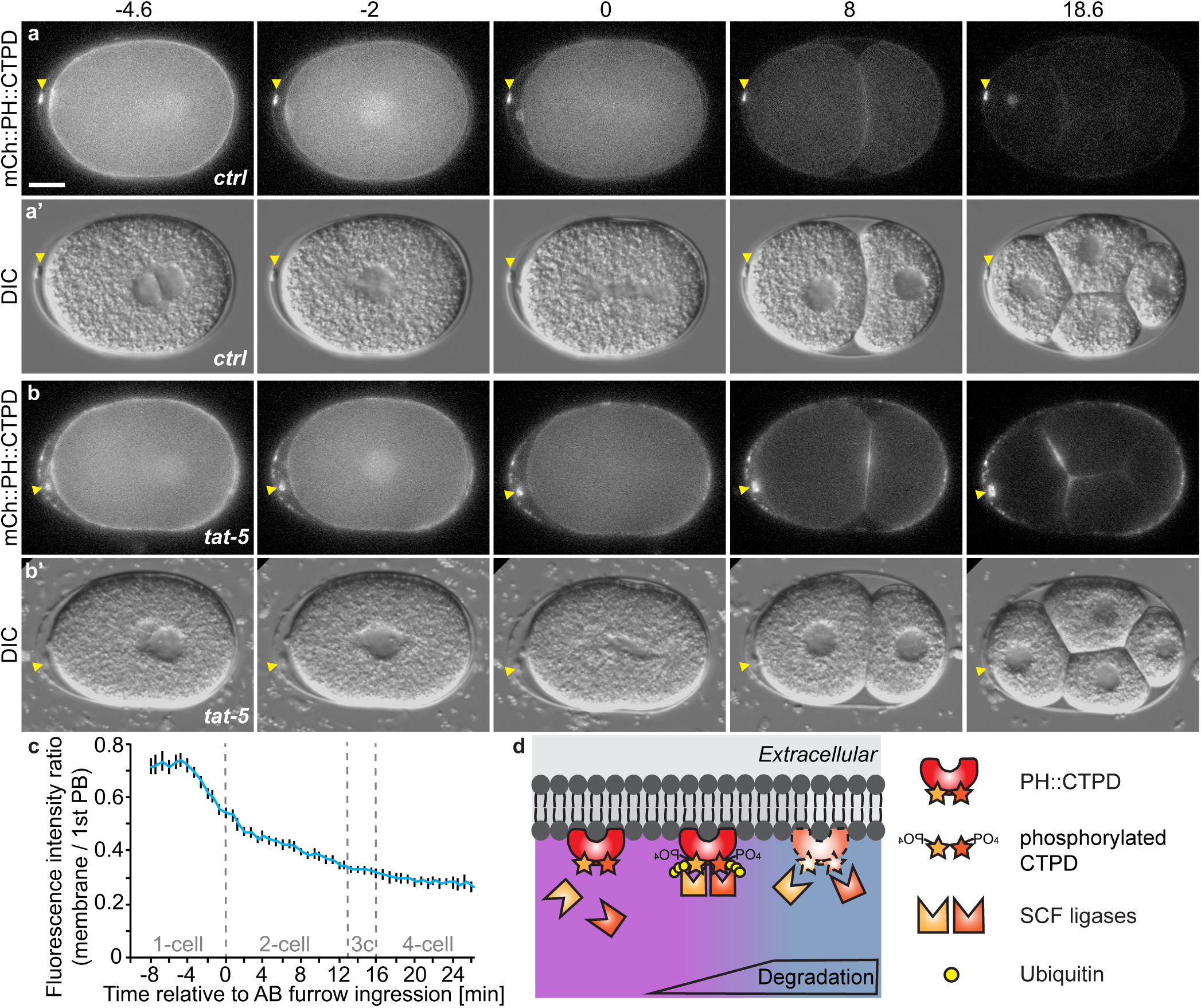
The C-terminal phosphodegrons (CTPD) of OMA-1 lead to transient degradation of a membrane reporter that preferentially labels extracellular vesicles. a) An mCherry and CTPD degron-tagged PH reporter localizes to the plasma membrane in 1-cell embryos, but begins to be degraded during mitosis. Weaker fluorescence persists in embryonic cells from the 2-cell stage on, but polar bodies remain brightly labelled (arrowhead, n=42). Scale bar: 10 µm. b) After *tat-5* knockdown, the mCh::PH::CTPD reporter still degrades, but bright patches of extracellular vesicles are visible in the egg shell, floating around the embryo, and on the embryo surface, in addition to the brightly labelled polar bodies (arrowhead, n=43). See also Video 2. c) Loss of mCh::PH::CTPD fluorescence from the cell surface begins ~3 minutes before cytokinetic furrow ingression, but degradation tapers off leaving residual mCh::PH::CTPD fluorescence after the 2-cell stage. Fluorescence intensity was normalized to the 1^st^ polar body to correct for photobleaching. Bars represent mean ± SEM. d) Phosphorylation of the CTPD degrons leads to recognition of the CTPD by multiple SCF ubiquitin ligases, ubiquitination, and degradation.

We also tested whether it was possible to label EVs by degron-tagging transmembrane proteins. In contrast to the proteasomal degradation of cytosolic proteins, ubiquitination of transmembrane proteins leads to endocytosis and lysosomal degradation^12^. The syntaxin SYX-4 is a single-pass transmembrane protein that localizes to the plasma membrane and endocytic vesicles (Fig. S2a-c)^22^. We tagged the cytosolic domain of SYX-4 with the ZF1 degron to make it accessible to ZIF-1. Degron-tagged GFP::ZF1::SYX-4 localizes normally before the onset of ZIF-1 expression (Fig. S2d), after which GFP::ZF1::SYX-4 accumulates in intracellular vesicles and is lost from the plasma membrane (Fig. S2e). These vesicles eventually disappear from ZIF-1-expressing cells (Fig. S2f), consistent with ubiquitin-driven endocytosis and lysosomal degradation (Fig. S2i). To test whether lysosomal degradation of a transmembrane protein can label EVs, we treated the GFP::ZF1::SYX-4 reporter strain with *tat-5* RNAi to induce microvesicle release. Similar to the degron-tagged PH reporter (Fig. 2k-n), GFP::ZF1::SYX-4 accumulates around cells after *tat-5* knockdown (Fig. S2g-h)^21^. Thus, both membrane-associated and transmembrane proteins can be tagged with degrons to specifically label EVs.

### Degron protection assay reveals topology of membrane-associated proteins

We next tested whether degron tagging could reveal insights into protein topology. Clathrin is enriched at the cell surface after *tat-5* knockdown^21^, but it was unclear whether this was due to increased clathrin inside the plasma membrane or due to the release of clathrin in EVs that accumulated next to the plasma membrane (Fig. 4f). Both possibilities were plausible, as clathrin-binding proteins are increased at the plasma membrane after *tat-5* knockdown, including ESCRT proteins^21^, and as clathrin was found in purified *Drosophila* and mammalian EVs^23,24^. Therefore, we asked whether a degron-tagged clathrin heavy chain reporter would be protected from degradation inside EVs or be accessible to degradation at the cell cortex.

**Fig. 4:**
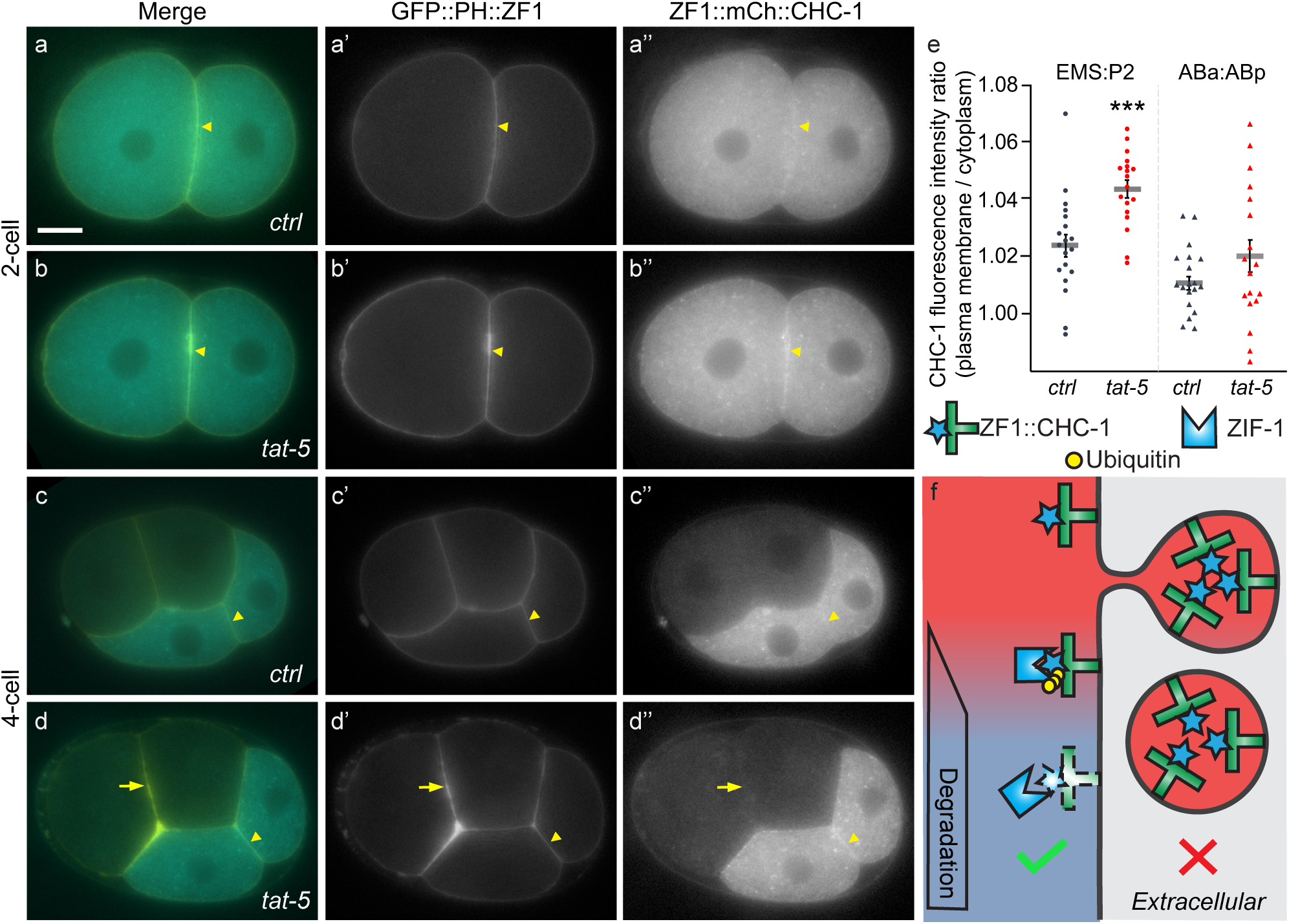
Degron protection assay reveals protein topology. a) A degron-tagged clathrin reporter ZF1::mCh::CHC-1 initially localizes to the plasma membrane (arrowhead) and intracellular puncta (n=9). Scale bar:10 µm. b) After *tat-5* knockdown, ZF1::mCh::CHC-1 is enriched at the plasma membrane (arrowhead, n=14). c) After expression of ZIF-1 begins in anterior cells, ZF1::mCh::CHC-1 is degraded throughout the anterior cells in control embryos (n=19). d) Although ZF1::mCh::CHC-1 is still enriched at the plasma membrane in posterior cells (arrowhead) in *tat-5* RNAi-treated embryos, ZF1::mCh::CHC-1 is lost from the plasma membrane in anterior cells (arrow, n=18), indicating that clathrin is accessible to ubiquitin ligases. GFP::PH::ZF1 labels the plasma membrane and extracellular vesicles. e) Quantification of clathrin enrichment on a posterior cell contact (EMS:P2) or anterior cell contact (ABa:ABp) compared to the neighbouring cytoplasm at the 4- and 6-cell stage from two independent experiments. ZF1::mCh::CHC-1 fluorescence was significantly increased at the posterior EMS:P2 cell contact (***p<0.001 using Student’s t-test, *ctrl* n=19, *tat-5* RNAi n=18). No change was observed at anterior cell contacts (p>0.05). Bars represent mean ± SEM. f) If clathrin were in extracellular vesicles, ZF1::mCh::CHC-1 would be protected from ZIF-1-mediated degradation. However, as clathrin is inside the plasma membrane, ZF1::mCh::CHC-1 is accessible to ZIF-1-mediated degradation.

Consistent with previous results^21^, we initially saw increased ZF1::mCh::CHC-1 and GFP::ZF1::PH labelling clathrin and membrane at cell contacts in embryos treated with *tat-5* RNAi (Fig. 4b) compared to control embryos (Fig. 4a). This increase persisted in posterior *tat-5* knockdown cells (Fig. 4e), but after ZIF-1 expression began in anterior cells, ZF1::mCh::CHC-1 disappeared from the cell surface in both control and *tat-5* RNAi-treated embryos (Fig. 4c-e), while GFP::ZF1::PH persisted in EVs after *tat-5* knockdown (Fig. 4d). This demonstrates that the increased clathrin signal is due to association with the plasma membrane and not due to clathrin trapped within EVs. Thus, degron-tagged reporters reveal whether a protein is inside or outside the plasma membrane.

### Degron reporters enable tracking of phagocytosed cargo

To test whether the specific labelling by degron tags facilitates long-term tracking, we observed the two polar bodies in which ZF1 degradation does not occur (Fig. 2d-f). As polar bodies are dying cells, they have a nucleus and can be tagged with chromosome reporters like histone H2B, in addition to the PH domain^20^. Both polar bodies are initially found on the anterior surface of the embryo (Fig. 5a, d). The first polar body is trapped in the eggshell^25^, while the second polar body (2PB) is phagocytosed by an anterior cell^20^. Because mCh::H2B labels all nuclei in the embryo, it can be challenging to track the 2PB phagosome among the many dividing nuclei (Fig. 5b-c, Video 3). In contrast, the degron-tagged ZF1::mCh::H2B reporter disappears sequentially from somatic nuclei, leaving the two polar bodies as the only fluorescent structures on the anterior half of the embryo (Fig. 5e, Video 3). This confirms that the intervening cell corpse and phagosome membranes protect ZF1-tagged proteins from proteasomal degradation (Fig. 5g). Improving the signal-to-noise ratio with degron-tagged reporters also improves automated tracking of the 2PB by removing overlapping traces (Fig. 5c, f). Tubulation of the 2PB phagosome into small vesicles can also be followed with degron-tagged reporters (video 1, left)^20^. Thus, degron-tagged reporters reveal organelle dynamics and facilitate tracking by removing background labelling.

**Fig. 5:**
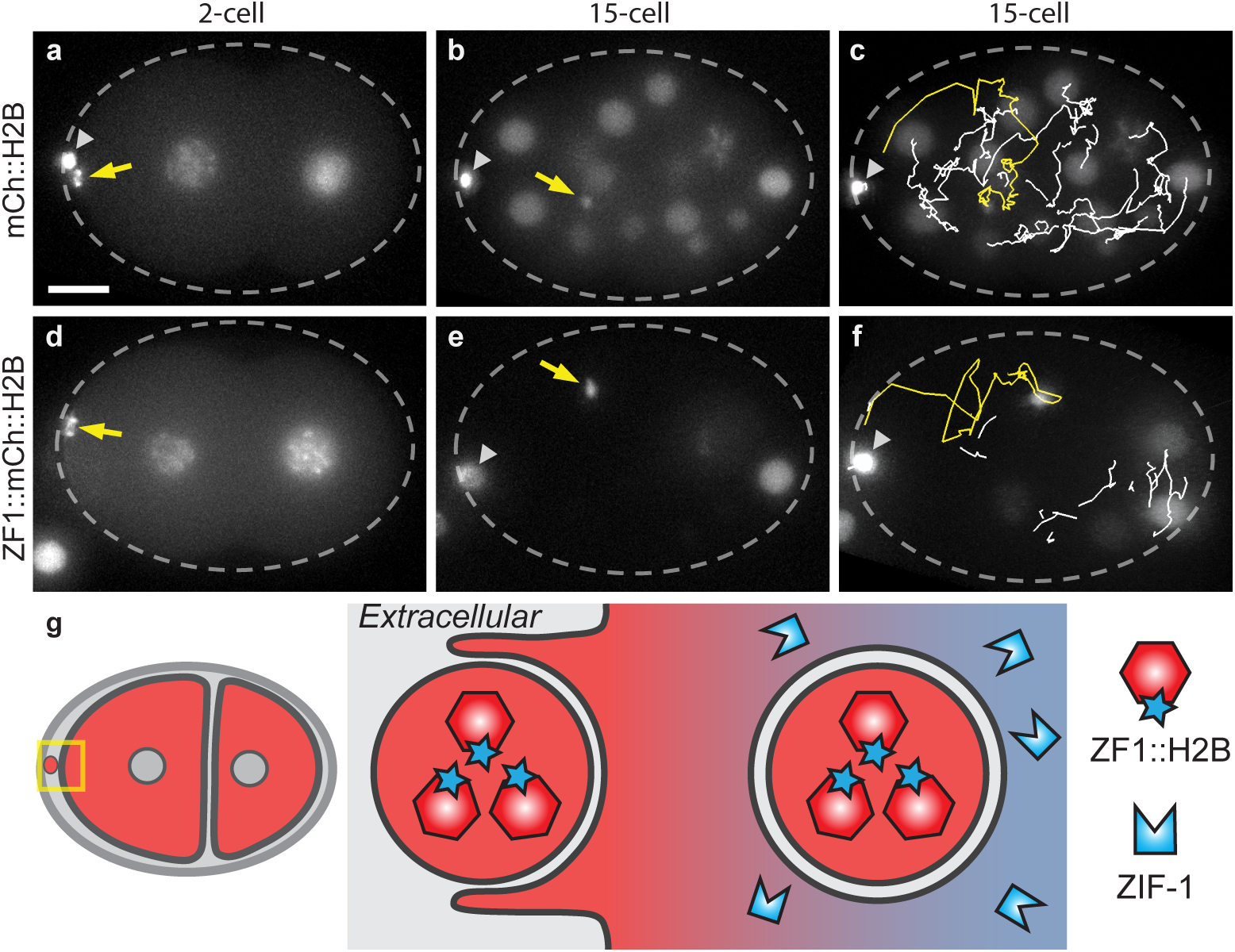
Degron reporters label phagocytosed cell debris and enable tracking. a-c) The histone reporter mCh::H2B labels chromosomes in embryonic nuclei and two polar bodies (n=13). The first polar body (arrowhead) is trapped in the eggshell (dashed oval). Scale bar: 10 µm. a) The second polar body (2PB, arrow) neighbours the anterior blastomere (AB cell) in a 2-cell embryo. b) The 2PB is engulfed in a phagosome in a 15-cell embryo. Due to H2B fluorescence from surrounding embryonic nuclei, it is hard to track the 2PB. c) Automated tracking of the 2PB (yellow) results in crossing tracks from nearby nuclei (white), increasing the potential need for manual correction. d-f) Embryos labelled with ZF1-tagged mCh::H2B (n=12). d) Released 2PB neighbours a 2-cell embryo. e) After engulfment, 2PB is easily trackable due to degradation of ZF1::mCh::H2B in somatic cell nuclei. See also video 3. f) Automated tracking of the 2PB (yellow) is simplified by removing the label of nearby nuclei (white). g) Degron reporters released in cells or other debris prior to expression of the ZIF-1 ubiquitin ligase adaptor are protected from ZF1-mediated degradation. After engulfment in a second layer of membrane, they are still protected from proteasomal degradation. As degron reporters are degraded in the cytosol, only the reporters within the phagosome remain fluorescent, improving the signal-to-noise ratio.

### Degron reporters reveal plasma membrane topology and dynamics

Degron-tagged reporters also allow the specific labelling of a single membrane or surface of a membrane. The close proximity of the corpse plasma membrane and the engulfing phagosome membrane make it difficult to distinguish their signals using fluorescence microscopy^26,27^, unless cells have distinct transcriptional programs, i.e. neuronal corpses engulfed by non-neuronal cells. Using degron reporters, this can be achieved even with sister cells *in vivo* by degrading the reporter localized to the phagosome membrane, leaving the corpse membrane preferentially labelled (Fig. 6d). For example, as polar bodies do not enter mitosis, their plasma membranes remain brightly labelled after mCh::PH::CTPD degradation occurs in the embryo (Fig. 6a). After phagocytosis, the 2PB membrane appears as a bright hollow sphere in a weakly labelled cell using the mCh::PH::CTPD reporter (Fig. 6b). In order to degrade or recycle corpse contents, the plasma membrane of engulfed corpses must be disrupted within the safety of the phagosome membrane^28^. 2PB membrane breakdown can be visualized using the mCh::PH::CTPD reporter, which is seen dispersing throughout the phagosome lumen (Fig. 6c). Similar results were found using the mCh::PH::ZF1 reporter (Video 1)^20^. Thus, by depleting fluorescent reporters from membrane surfaces facing the cytosol, degron-tagging enables examination of specific membranes and their dynamics.

**Fig. 6:**
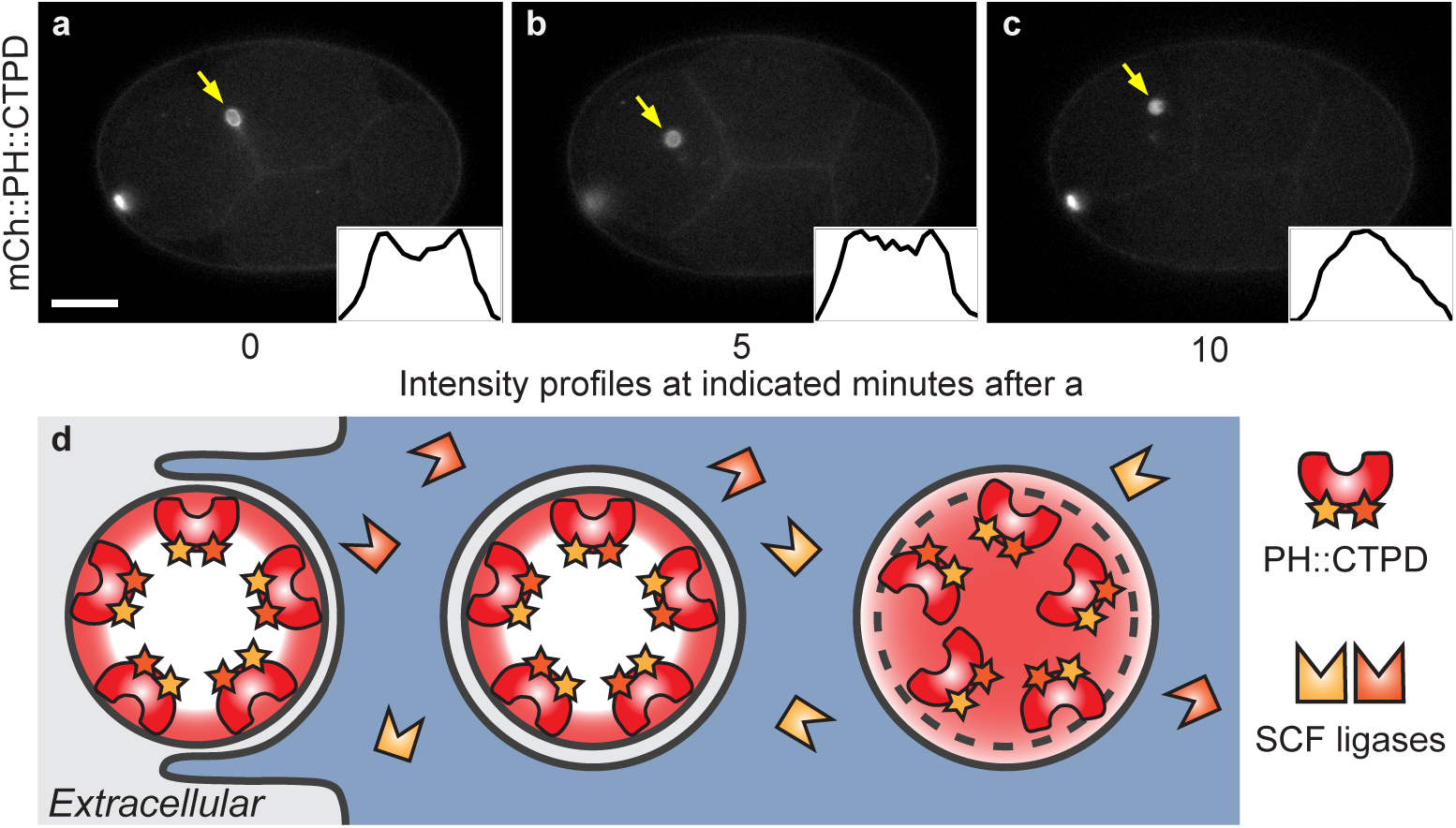
Degron reporters label specific membranes. a) The mCh::PH::CTPD reporter brightly labels the plasma membrane of the polar bodies and weakly labels the plasma membrane of embryonic cells at the 4-cell stage (n=22). Scale bar: 10 µm. b) After phagocytosis of the second polar body (2PB, arrow), the plasma membrane of the 2PB is distinctly visible. c) Several minutes later, the corpse plasma membrane breaks down inside the phagolysosome and the mCh::PH::CTPD reporter disperses throughout the phagolysosome lumen, as demonstrated by line scans through the phagolysosome (insets). See also video 2. d) The mCh::PH::CTPD reporters inside the corpse membrane are protected from degradation. Model of membrane breakdown inside the phagolysosome, demonstrating the shape change from a hollow to a filled sphere.

### Degron protection assays reveal nuclear membrane topology and dynamics

We next asked whether degron-tagged reporters can be used to assess nuclear topology. Other components of the ECS ubiquitin ligase complex are found in both the cytosol and nucleus, but the ZIF-1 ligase adaptor appears cytosolic^29^. To test whether ZIF-1 degradation was restricted to the cytosol (Fig. 7a), we tagged the nuclear lamin LMN-1 with the ZF1 degron to examine the dynamics of the nuclear cortex during cell division^30^. Prior to the onset of ZIF-1 expression, the fluorescence intensity of the mKate2::ZF1::LMN-1 reporter is comparable between the anterior and posterior nuclei (Fig. 7c). After ZIF-1 expression begins in the two anterior daughter cells, the interphase levels of mKate2::ZF1::LMN-1 gradually drop in comparison to posterior cells (Fig. 7d, g), suggesting that some degradation occurs despite the intact nuclear envelope. However, the degradation of the mKate2::ZF1::LMN-1 occurred faster during mitosis (Fig. 7e-g, video 4). The nuclear envelope breaks down (NEBD) during mitosis for chromosome segregation^31^, which would allow cytosolic ZIF-1 to target ZF1-tagged nuclear proteins (Fig. 7b).

**Fig. 7:**
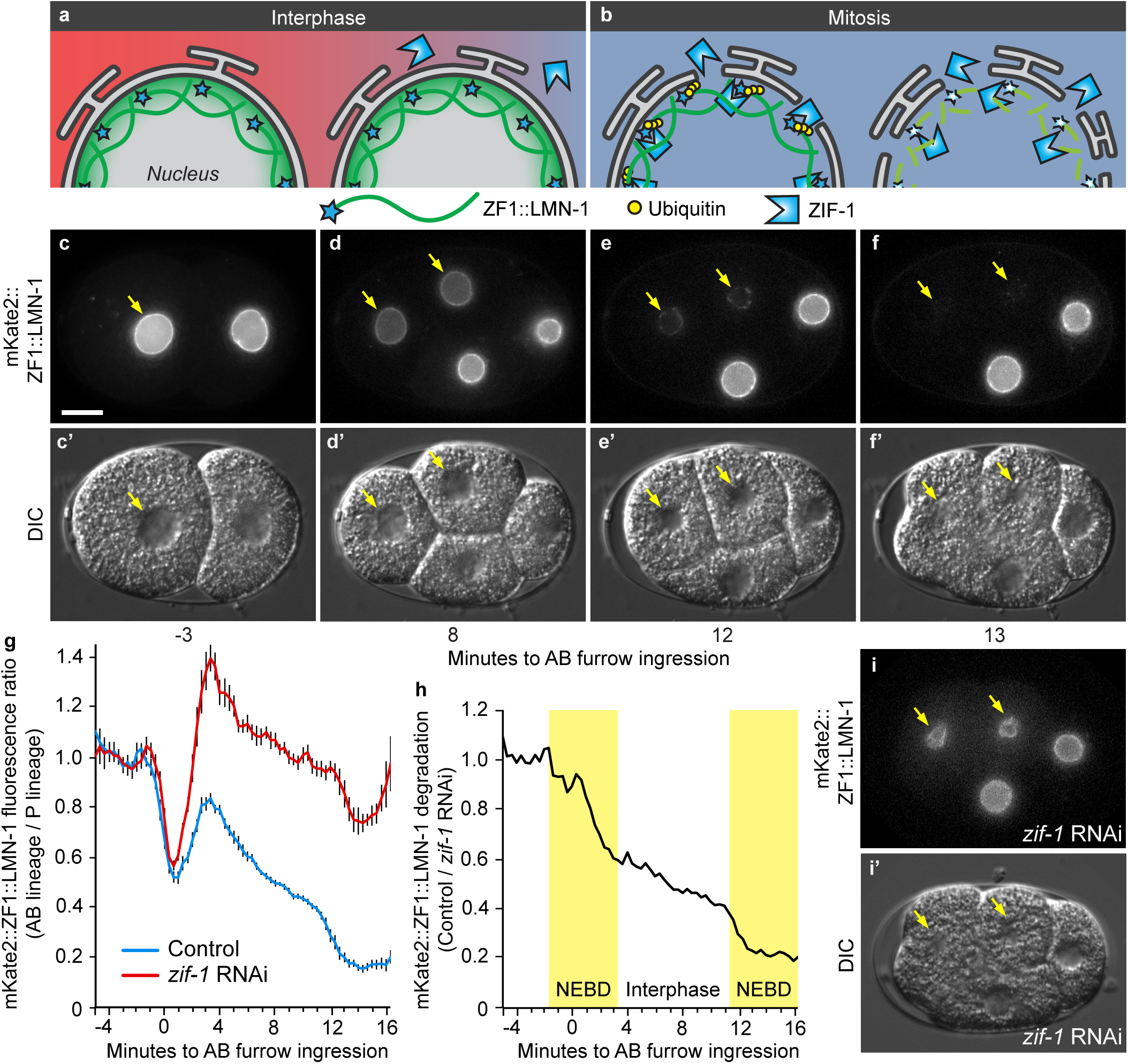
Degron reporters reveal nuclear envelope topology during cell division. a) During interphase, the nuclear envelope is intact, hindering ZIF-1 binding to ZF1-tagged reporters inside the nuclear envelope. b) During mitosis, the nuclear envelope is remodelled into the endoplasmic reticulum, allowing binding of ZIF-1 to ZF1-tagged reporters, which leads to ubiquitination and rapid proteasomal degradation. c) Prior to ZIF-1 expression, mKate2::ZF1::LMN-1 is similarly bright in the nuclear matrix of the anterior AB cell (arrow) and the posterior cell (n=14). Scale bar: 10 µm. d) After the onset of ZIF-1 expression in anterior daughter cells (ABx), mKate2::ZF1::LMN-1 starts to lose fluorescence in anterior cells (arrows). e-f) During mitosis, the nuclear envelope is disassembled, leading to rapid proteasomal degradation of mKate2::ZF1::LMN-1. c’-f’) DIC images show nuclear morphology during cell cycle progression. g) Quantification of mKate2::ZF1::LMN-1 fluorescence in the dorsal ABp nucleus compared to the ventral EMS nucleus with (n=5-11) and without *zif-1* RNAi treatment (n=5-14). Bars represent mean ± SEM. h) Normalizing mKate2::ZF1::LMN-1 to *zif-1* RNAi-treated embryos demonstrates that fluorescence drops 4-6 times faster during nuclear envelope breakdown (NEBD) in anterior cells (yellow boxes, AB: 16% per min, ABx: 11% per min), when ZIF-1 has full access to mKate2::ZF1::LMN-1. Slow decline in the intensity of mKate2::ZF1::LMN-1 is also visible during interphase (3% per min). NEBD was defined as the first time point the nuclear lamina collapsed in the mother cell until the lamina became round again in daughter cells. Time is given in relation to AB furrow ingression. i) mKate2::ZF1::LMN-1 is maintained in anterior cells after *zif-1* knockdown (n=11). i’) DIC image showing similar stage to f. See also video 4.

To confirm that loss of fluorescence was due to ZIF-1-mediated degradation and not due to morphological changes in the nuclear envelope during the cell cycle, we treated reporter embryos with *zif-1* RNAi to inhibit degradation. The fluorescence of mKate2::ZF1::LMN-1 fluctuated as cells divided (Fig. 7g), but fluorescence persisted in anterior cells (Fig. 7i). We normalized the control curve to the *zif-1* curve to remove changes due to nuclear morphology, revealing 4-6 times faster degradation during NEBD (Fig. 7h). Thus, although a pool of mKate2::ZF1::LMN-1 is accessible to ZIF-1 during interphase, mKate2::ZF1::LMN-1 is largely protected from proteasomal degradation by the nuclear envelope (Fig. 7a). After treating mKate2::ZF1::LMN-1 embryos with *zif-1* RNAi, we noticed an increase in nuclear morphology defects (Fig. S1c). However, the mKate2::ZF1::LMN-1 reporter did not result in other defects typical of LMN-1 overexpression, even when *zif-1* was knocked down (Fig. S1d-f). Therefore, degron-tagging provides probes for membrane topology that can investigate the dynamics of nuclear envelope breakdown.

As LMN-1 is largely immobile in the nuclear lamina^32^, the observation that some degradation occurred during interphase raised the possibility that ubiquitination of the degron-tagged reporter was altering LMN-1 dynamics. To determine whether mKate2::ZF1::LMN-1 became mobile after ZIF-1 expression, we performed fluorescence recovery after photobleaching (FRAP) experiments in an anterior daughter nucleus (Fig. S3a). There was no significant recovery of mKate2::ZF1::LMN-1 fluorescence during interphase (Fig. S3b), suggesting that the mobility of LMN-1 was unchanged by ubiquitination. Thus, quantification of degron-tagged reporters can be a tool to examine protein dynamics in addition to organelle dynamics.

### The localization of ubiquitin ligase adaptors determines the localization of degradation

We next tested whether protection by internal membranes was a specific feature of ZIF-1 and ECS ubiquitin ligases or whether the nuclear envelope could also protect targets of SCF ubiquitin ligases from degradation. As SCF ubiquitin ligase complexes are found in the nucleus and cytosol, we modified the TIR1 ligase adaptor from plants with a nuclear export signal (NES) and expressed NES-TIR1 in HeLa cells that also expressed a lamin A reporter tagged with a minimized AID (mAID) degron^33^. During interphase, little Venus-mAID-LMNA fluorescence was lost when TIR1 was restricted to the cytosol (Fig. 8a, c, f, Video 5). After cells entered mitosis and the nuclear envelope broke down (NEBD), NES-TIR1 caused rapid degradation of Venus-mAID-LMNA (Fig. 8b-c, f, Video 6). Thus, degron protection assays can also be used in mammalian cells.

**Fig. 8:**
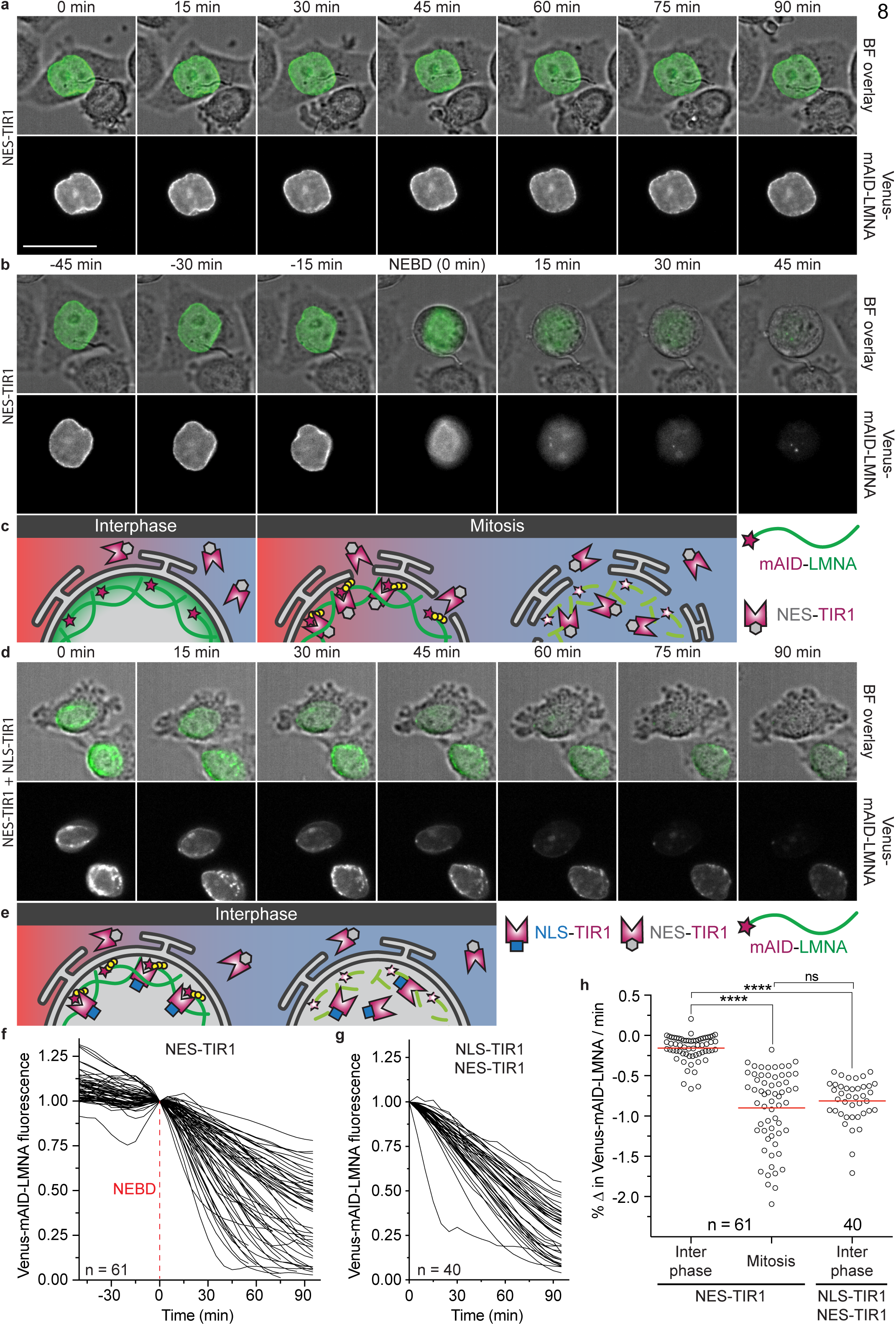
Localization of the ubiquitin ligase adaptor spatially controls degradation. a) Representative time-lapse images of a HeLa cell transiently expressing NES-TIR1 and Venus-mAID-LMNA. 0.5 mM NAA was added at t = 0 to induce auxin-dependent degradation. During interphase, Venus-mAID-LMNA in the nuclear lamina is protected from ubiquitination by cytosolic NES-TIR1 and fluorescence persists in the presence of NAA. Scale bar: 30 µm. b) The same cell is shown entering mitosis. After nuclear envelope breakdown (NEBD, t = 0), Venus-mAID-LMNA is accessible by cytosolic NES-TIR1 and fluorescence rapidly disappears. c) Model of mAID-LMNA protection from cytosolic NES-TIR1 during interphase and accessibility to cytosolic NES-TIR1 after NEBD occurs during mitosis. d) Interphase HeLa cell stably expressing NLS-TIR1 and NES-TIR1 from a single mRNA in addition to Venus-mAID-LMNA. After treatment with 0.5 mM NAA (t = 0), Venus-mAID-LMNA fluorescence rapidly disappears during interphase. e) Model of mAID-LMNA accessibility to nuclear NLS-TIR1 during interphase. f) Quantification of Venus-mAID-LMNA fluorescence before and after NEBD (t = 0) in the presence of cytosolic NES-TIR1 and NAA. Values were normalized to NEBD. g) Quantification of Venus-mAID-LMNA fluorescence in interphase cells in the presence of cytosolic NES-TIR1, nuclear NLS-TIR1, and NAA. Values were normalized to t = 0. h) Comparison of Venus-mAID-LMNA degradation velocities from single cells. The degradation velocity was determined for 45 minutes before NEBD (interphase) and 45 minutes after NEBD (mitosis) for NES-TIR1-expressing cells from three independent experiments or during the first 45 minutes after NAA addition for cells expressing both NLS-TIR1 and NES-TIR1 from two independent experiments. Bars indicate the mean of the indicated number of cells. Significance according to unpaired multi-comparison Kruskal-Wallis test with Dunn’s statistical hypothesis testing (****, p<0.0001; ns, p>0.9999).

To confirm that protection from degradation was due to the localization of the ligase adaptor and not due to a protective effect of the nuclear lamina, we co-expressed NLS-TIR1, which has a nuclear localization signal (NLS), with NES-TIR1^33^. When TIR1 was localized to both the nucleus and the cytosol, Venus-mAID-LMNA was rapidly degraded during interphase (Fig. 8d-e, g, Video 7). The velocity of interphase degradation by NLS-TIR1 was not significantly different from mitotic degradation by NES-TIR1 (Fig. 8h), indicating that LMNA was not protected from ubiquitination by integration into the nuclear lamina. These results demonstrate that the localization of the ligase adaptor can determine the localization of degradation, providing a strategy to target one pool of a reporter protein for degradation.

### Degron reporters reveal membrane topology during abscission

In order to understand how degrons in restricted spaces can be ubiquitinated and degraded, we applied degron-tagged reporters to the process of abscission. During cell division, the actomyosin furrow closes around the spindle midbody to form a narrow intercellular bridge^34^. Both sides of the bridge are cleaved during abscission to release a ~1 µm extracellular vesicle called the midbody remnant, which is later phagocytosed (Fig. 9h)^35^. The intercellular bridge no longer permits diffusion between cells ~4 minutes after furrow ingression^34^, but the first cut for abscission does not occur until ~10 minutes after furrow ingression^36^. Therefore, we asked when actomyosin accumulated in the intercellular bridge was accessible for proteasomal degradation. We tagged non-muscle myosin (NMY-2) with the ZF1 degron and measured the fluorescence intensity of NMY-2::GFP::ZF1 in the bridge between the anterior daughter cells (Fig. 9a)^35^. Degradation of cytoplasmic NMY-2::GFP::ZF1 was first visible 8 ± 1 minutes after furrow ingression. NMY-2::GFP::ZF1 in the bridge showed a small but significant decline for the next 2 minutes (Fig. 9g), suggesting that NMY-2 is normally able to diffuse out of the bridge up to 10 minutes after furrow ingression. Subsequently, NMY-2::GFP::ZF1 in the bridge was protected from proteasomal degradation (Fig. 9b, g), suggesting that either a diffusion barrier had formed or abscission had occurred. Thus, using degron reporters and light microscopy on living embryos, we could confirm the timing of abscission estimated from electron microscopy data from fixed embryos.

**Fig. 9:**
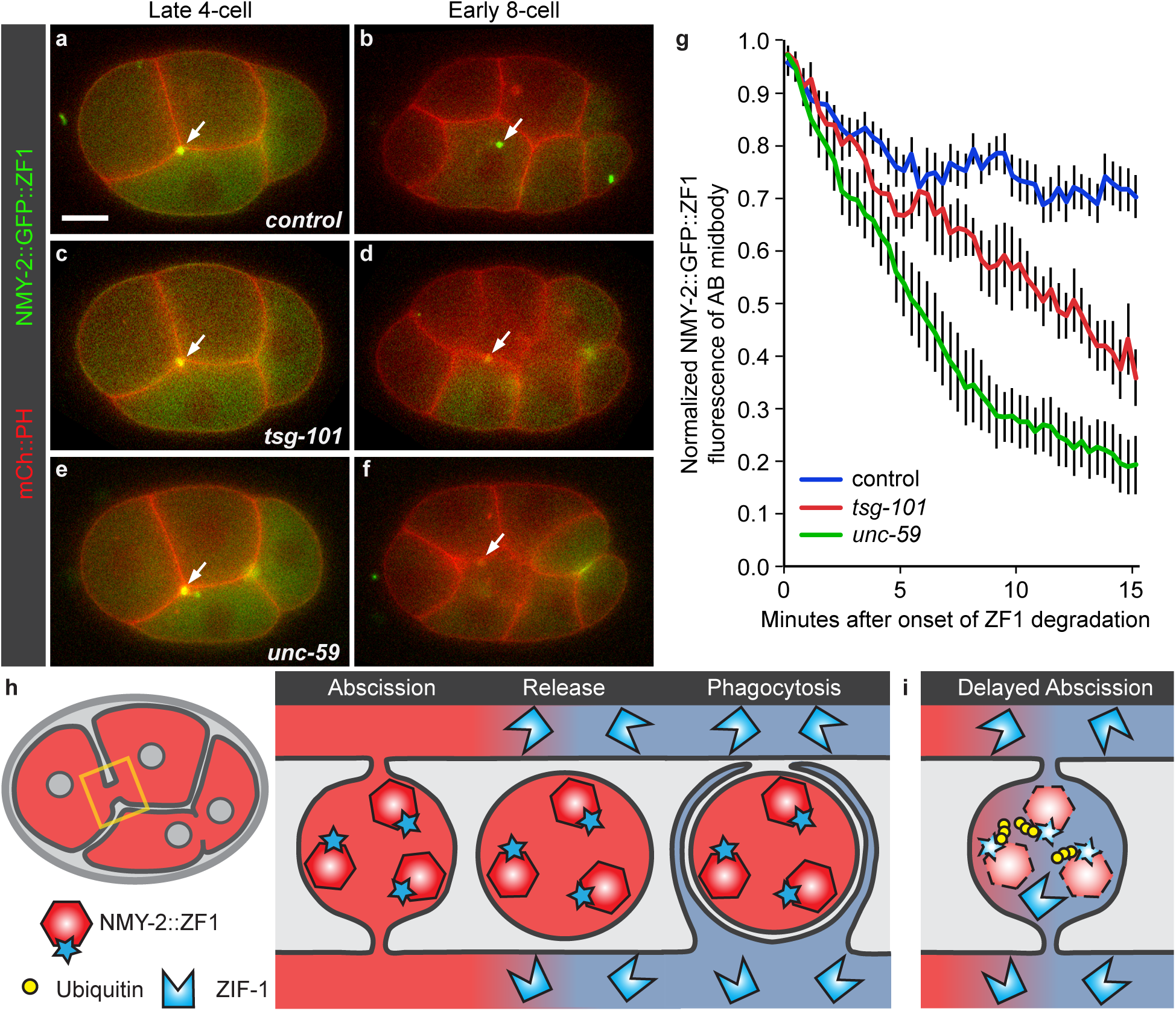
Degron reporters reveal membrane topology during abscission. a-f) Time-lapse images of embryos expressing mCh::PH to label the plasma membrane and NMY-2::GFP::ZF1 to label non-muscle myosin in the cytokinetic ring. Scale bar: 10 µm. a) In a control embryo, NMY-2::GFP::ZF1 between the anterior daughter cells (arrow) is protected from ZF1-mediated degradation due its release outside cells in the midbody remnant after abscission. b) NMY-2::GFP::ZF1 in the phagocytosed midbody remnant is protected from proteasomal degradation by engulfing membranes (n=19). c) In embryos depleted of the ESCRT-I subunit TSG-101, NMY-2::GFP::ZF1 localizes to the intercellular bridge normally. d) NMY-2::GFP::ZF1 is degraded after *tsg-101* knockdown, indicating that abscission is incomplete and NMY-2::GFP::ZF1 is accessible to the degradation machinery. Engulfment of the AB midbody is also delayed, likely due to incomplete abscission (n=7). e-f) Embryos depleted of the septin UNC-59 show rapid degradation of NMY-2::GFP::ZF1 from the intercellular bridge and delayed engulfment of the midbody remnant due to defects in abscission (n=8). g) Fluorescence intensity of the NMY-2::GFP::ZF1 reporter on the intercellular bridge between anterior daughter cells (AB midbody). NMY-2::GFP::ZF1 fluorescence in the bridge drops significantly for two minutes after the onset of ZF1 degradation in the cytoplasm of control embryos (blue, n=11, p<0.05 using Student’s t-test with Bonferroni correction), showing that NMY-2 is able to diffuse out of the bridge. Fluorescence then persists, showing that formation of the diffusion barrier, symmetric abscission, and engulfment protect the degron-tagged reporter from degradation. In contrast, NMY-2::GFP::ZF1 fluorescence continues to drop in *tsg-101* RNAi-treated embryos (red, n=5, p<0.05 compared to control after 8 min) and *unc-59* mutants (green, n=6, p<0.05 compared to control after 3.5 min), showing that the ZF1 degron technique is sensitive to detecting small defects in abscission. Bars represent mean ± SEM. h) In control embryos, degron reporters in the intercellular bridge are protected from proteasomal degradation due to release after abscission and encapsulation in a phagosome. i) In abscission mutants, degron reporters are accessible to the cytosol, leading to their removal by proteasomal degradation.

To test whether degron-tagged reporters were able to detect novel phenotypes in abscission mutants, we depleted proteins implicated in abscission, including the ESCRT-I subunit TSG-101 and the septin UNC-59. At the onset of ZF1-mediated degradation, NMY-2::GFP::ZF1 labelled intercellular bridges in *tsg-101* or *unc-59* mutants normally (Fig. 9c, e). In contrast to control embryos where NMY-2::GFP::ZF1 fluorescence persisted through midbody release and phagocytosis (Fig. 9b), NMY-2::GFP::ZF1 fluorescence intensity continued to drop significantly in the bridge of both *tsg-101* and *unc-59* mutants (Fig. 9d, f-g), and this drop was dependent on ZIF-1 expression^35^. Phagocytosis of the midbody remnant was also delayed in both *tsg-101* and *unc-59* mutants (Fig. 9d, f)^34,35^, consistent with a delay in abscission. These findings demonstrate that NMY-2 is able to diffuse out of the bridge and be degraded when abscission is delayed (Fig. 9i). No defect was detected for *tsg-101* knockdown using a dextran diffusion assay^34^, demonstrating the high sensitivity of the degron protection assay for detecting abscission defects.

### Single membrane bilayers are sufficient to protect proteins from degron-mediated degradation

As all of the model systems we examined involved two membrane bilayers between the ubiquitin ligase and the degron-tagged reporter (nucleus, extracellular vesicle, phagocytosed cell debris), we tested whether a single membrane bilayer would protect a reporter from degradation. We expressed a degron-tagged reporter in the secretory pathway and maintained it in the endoplasmic reticulum (ER) using a C-terminal KDEL sequence^37^ (Fig. 10d). The ss::TagRFP-T::ZF1::KDEL reporter localized to the ER, similar to established reporters (Fig. 10a-b). After ZIF-1 expression began in anterior cells, ss::TagRFP-T::ZF1::KDEL persisted with no measurable loss of fluorescence (Fig. 10b-c). Thus, a single membrane bilayer protects degron-tagged reporters from degradation, confirming that degron protection assays can be applied to intracellular organelles.

**Fig. 10:**
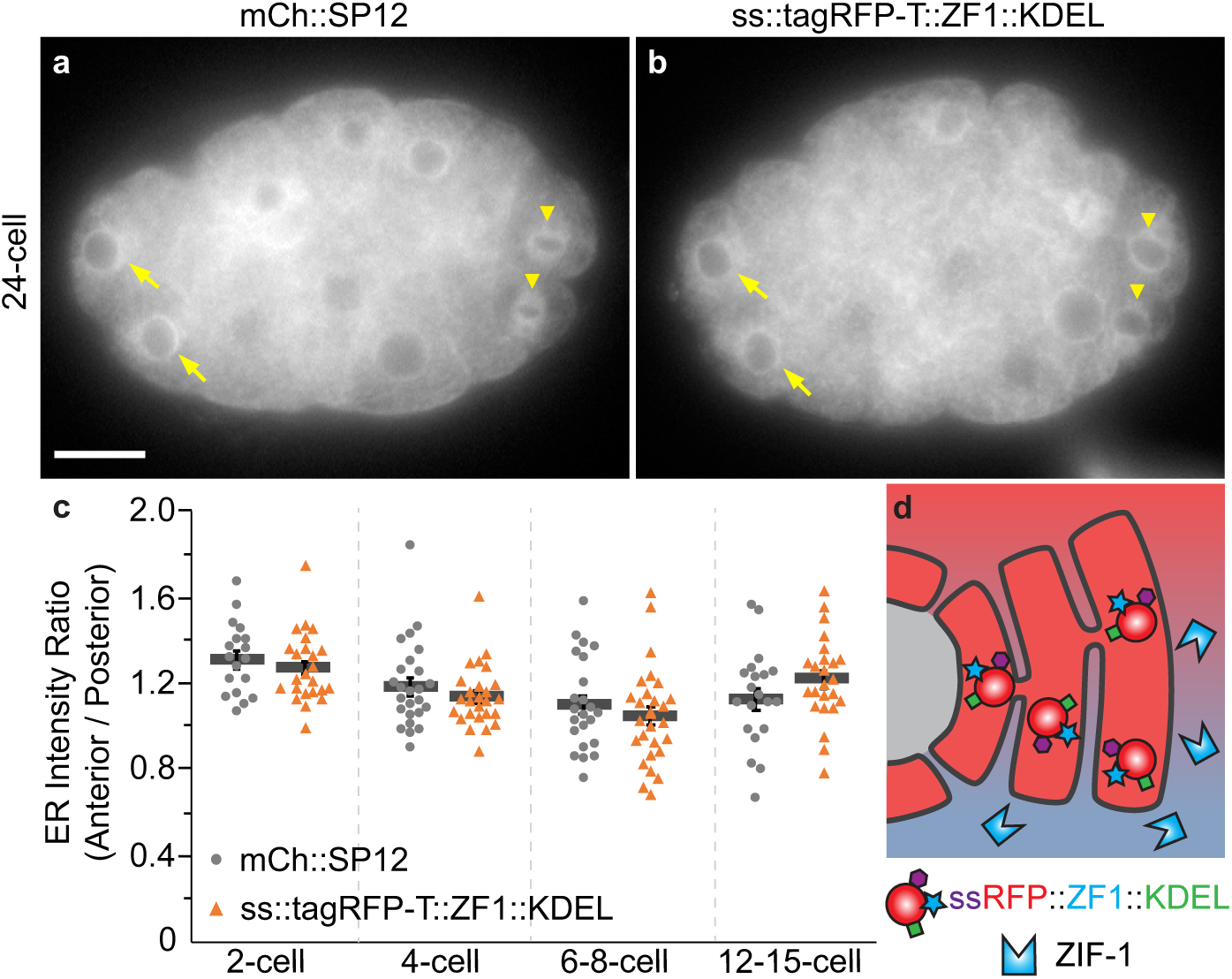
A ZF1 degron reporter in the ER is protected from ZIF-1/ECS-mediated degradation. a) mCherry-tagged SP12 localizes to the ER similarly in anterior (somatic, arrow) and posterior (germ line, arrowhead) cells (n=102). Scale bar: 10 µm. b) An RFP- and ZF1 degron-tagged KDEL reporter similarly localizes to the ER in both anterior (somatic, arrow) and posterior (germ line, arrowhead) cells, despite expression of the ZIF1 ligase adaptor in anterior cells (n=88). c) There was no significant difference in the ratio between the fluorescence of anterior AB lineage cells and of posterior germ line or germ line sister cells at the indicated stages (p>0.05, Student’s t-test with Bonferroni correction mCh::SP12 n=18-25, ss::tagRFP-T::ZF1::KDEL n=24-28). Bars represent mean ± SEM. d) The ER membrane protects luminal proteins from ubiquitination by cytosolic ligase complexes.

### Degron-tagged reporters did not result in degradation of binding partners

Ligase adaptors can target proteins for ubiquitination through intermediary binding partners, including nanobodies^33^, which raised the possibility that degron-tagged reporters would lead to the degradation of untagged binding partners. This could be especially relevant when the reporter protein assembles into larger complexes, such as histones or the nuclear lamina, or dimerizes, such as NMY-2. To test whether degron-tagged proteins lead to the degradation of untagged proteins, we generated strains expressing fluorescent reporter proteins for H2B, LMN-1, and NMY-2 both with and without the ZF1 degron tag. We measured the intensity of fluorescence for each reporter and found that although the ZF1-tagged reporter degraded, the reporter without the degron did not show significant changes in comparison to a strain that did not express any degron-tagged reporter (Fig. S4). Thus, degron-tagging does not generally lead to degradation of protein complexes.

## Discussion

In summary, degron-mediated degradation is more than a loss-of-function technique; it is a powerful tool to study dynamics from the level of proteins to organelles to cells. By removing cytoplasmic fluorescence, degron tags improve the visibility of extracellular, luminal, or nuclear reporters and enable long-term tracking. Degron-tagged reporters reveal insights on an epifluorescence microscope that are typically limited to super-resolution or electron microscopy on fixed samples. Our studies have focused on structures that are protected by membrane bilayers, but this approach should work for any structure resistant to ubiquitination or diffusion. Thus, degron-tagging is an important addition to the cell biologist’s toolbox.

As degron tags are widely used in cell extracts, cell culture and *in vivo*, this approach can visualize structures in many systems. We started with endogenous degrons in *C. elegans* embryos for their simplicity, only requiring expression of a degron-tagged reporter. Heterologous expression of ZIF-1 in worms or zebrafish can also degrade ZF1-tagged proteins and could be adapted to more systems^15,38^. Fusing an anti-GFP nanobody to endogenous ubiquitin ligase adaptors like ZIF-1 enables spatial control of degradation of GFP-tagged proteins in *C. elegans*, *Drosophila*, plants, and zebrafish^38–41^, which allows existing GFP-tagged reporters to be used for degron protection assays. Heterologous expression of the auxin-inducible degron (AID) and TIR1 ligase adaptor is also used in various animal models^18,33,42^. We used AID in mammalian cells to demonstrate that altering the localization of the ubiquitin ligase adaptor is sufficient to target degradation to the nucleoplasm or cytoplasm, consistent with observations on endogenous ligase adaptors^43^. Thus, the spatial control of degradation enabled by heterologous expression of ligase adaptors refers not only to expression in specific cell types, but also to specific organelles. As different ubiquitin ligases are found in different cellular compartments^44^, the choice of degron-ligase pair can enable degron protection assays in a range of organelles. For example, the Ab-SPOP/FP system uses a BCR ubiquitin ligase complex including Cul3 to specifically target GFP-tagged proteins in the nucleus for degradation^38,45^, but Cul3 also localizes to the Golgi. Furthermore, ubiquitin-mediated degradation can be regulated by small molecule drugs, temperature, or light^11^, offering many modalities to control the timing and localization of protein degradation. Thus, degron protection assays can readily probe cell biology in many model systems.

To avoid loss-of-function effects from induced degradation of degron-tagged proteins, our studies were performed with isolated protein domains or in the background of the untagged endogenous protein. Still, degradation of an overexpressed degron-tagged protein could alter some processes. We found that strains expressing degron-tagged fluorescent reporters were healthier than strains expressing fluorescent-tagged reporters. In fact, we only observed effects on nuclear morphology after inhibiting degron-mediated degradation of a LMN-1 reporter (Fig. S1c). Regardless, degrons like CTPD or AID that lead to partial or reversible degradation may be advantageous to avoid loss-of-function effects. In addition, it is important to verify that the degron tag does not alter the protein or process under study, similar to other protein tags. We did not observe changes to protein dynamics after degron tagging, with the expected exception of induced endocytosis of a transmembrane protein. Thus, degron tags are useful for probing the topology of transmembrane proteins, but not necessarily for studying their intracellular trafficking.

We used degron tags to label and track structures for which conventional reporters are insufficient. For example, extracellular vesicles (EV) are typically detected by the tetraspanin proteins on their surface, but tetraspanin content is heterogeneous among EV subpopulations^46^. By degron-tagging membrane-associated or transmembrane proteins, we were able to specifically label EVs *in vivo*, which has proven to be a valuable tool to screen for new regulators of EV budding^2^. Although our PH::ZF1 reporter is likely to favour plasma membrane-derived EVs (microvesicles), it is possible to target endosome-derived EVs (exosomes) by degron-tagging proteins associated with the endosome surface. Alternatively, exosomes may also be labelled by degron-tagging transmembrane proteins, given that ubiquitination drives endocytosis of transmembrane proteins (Fig. S2i). Degron-tagging reporters found at both the plasma membrane and endosome surface, such as actin-binding domains or abundant proteins like GAPDH, should label both microvesicles and exosomes^47^. Thus, degron-tagged reporters are uniquely able to specifically label EVs, enabling *in vivo* tracking and functional studies.

Wide-field light microscopy is normally limited to detecting structures that are >200 nm away from each other^48^. At this resolution, neighbouring membranes cannot be distinguished. In addition to specifically labelling extracellular vesicles next to the plasma membrane, we showed that degron-tagged reporters could distinguish the cargo corpse membrane from the engulfing phagosome membrane. This enabled the visualization of corpse membrane dynamics during phagolysosomal clearance using wide-field microscopy^20^. These reporters enable the precise staging of phagosomes for approaches such as correlative light and electron microscopy (CLEM)^49^, which can be used to determine the ultrastructure of membrane breakdown during phagolysosomal clearance. Therefore, degron-tagging is a useful tool to reveal novel insights into organelle dynamics in addition to the long-term tracking of specific cells *in vivo*.

Using a degron-tagged reporter, we were able to measure early changes in the topology of the nuclear envelope during cell division as well as in the restricted space of the intercellular bridge, which was previously only possible using electron microscopy^36^. We were also able to detect defects in abscission after knocking down an ESCRT subunit^35^, which were not visible using cytosolic diffusion assays^34^. Degron protection is therefore a sensitive tool to study the topology of membranes while they undergo fusion or fission.

Degron protection also revealed protein dynamics and topology. Knocking down septins or TSG-101 led to distinct rates of degron-mediated degradation of the myosin reporter, which could indicate that the intercellular bridge is open to differing degrees in these mutants, resulting in different rates of diffusion out of the bridge. We also found that a pool of the degron-tagged lamin reporter underwent degradation during interphase. This may indicate that an undetected level of ZIF-1 is in the nucleus, which ubiquitinates the ZF1 degron during interphase for degradation by the nuclear proteasome^50^. Alternatively, this may be due to undetected mobility of nuclear lamins into the cytosol during interphase. As cytosolic ligases are known to degrade proteins exported from the ER, while ER contents are protected from ECS- or SCF-mediated degradation (Fig. 10)^51^, degron-tagged reporters may also be used to reveal information on protein import/export across organelle membranes. As degradation of a reporter depends on its orientation as well as its location within the cell, degron protection assays can be applied to determining the topology of transmembrane proteins (type I/II) by testing whether they are accessible to degron-induced endocytosis and degradation, similar to our observations with SYX-4. Similarly, degron protection assays can distinguish cytosolic from luminal proteins. In summary, degron-tagged reporters improve the signal-to-noise ratio, reveal super-resolution insights on a standard microscope, and provide insights into localization and dynamics from the level of cells to proteins.

## Acknowledgements

Michaela Geisenhof and Alida Melse provided technical assistance. The imaging facility of the Rudolf Virchow Center and the Light Microscopy Facility of the BIOTEC at TU Dresden provided support for imaging and data analysis. Strains and reagents were provided by Peter Askjaer, Zhirong Bao, Michael Glotzer, Barth Grant, Jeremy Nance, Karen Oegema, Jonathon Pines, Christian Pohl, and the *Caenorhabditis* Genetics Center (CGC), which is funded by NIH Office of Research Infrastructure Programs (P40 OD010440). This work was funded by Deutsche Forschungsgemeinschaft (DFG) grants FA1046/3-1 to GF and WE5719/2-1 to AMW, and European Research Council (ERC) Horizon 2020 Research and Innovation Program grant 680042 to JM. We thank Avital Rodal, Cassie Blanchette, Katrin Heinze, Sonja Lorenz, and Anna Liess for critical reading of this manuscript.

## Author contributions

AMW and JM conceived, designed, and supervised the study. KBB, GF, KJ, LI, JC, and AMW performed and interpreted experiments. KBB, GF, and AMW wrote the manuscript.

## Materials and methods

### Worm strains and maintenance

*Caenorhabditis elegans* strains were maintained according to standard protocol at room temperature or 25°C^52^. For a list of strains used in this study, see Table S1. *unc-59* loss-of-function mutant embryos were generated by feeding *unc-59* RNAi to the WEH132 strain bearing a hypomorphic *unc-59* mutation^35^.

### Mammalian Cell Culture

Cells were cultured according to standard mammalian tissue culture protocols including testing for mycoplasma. HeLa Kyoto and HeLa FRT/TO cells were a kind gift from Jonathon Pines (ICR, London, UK). HeLa Kyoto or HeLa FRT/TO stably expressing NLS-TIR1_P2A_NES and Venus-mAID-LMNA were maintained in DMEM (Gibco) supplemented with 10% FBS, 1% (v/v) penicillin-streptomycin, 1% (v/v) Glutamax and 0.5 µg/ml Amphotericin B. A Neon Transfection system (Thermo Fisher Scientific) was used to generate HeLa FRT/TO NLS-TIR1_P2A_NES-TIR1 cells^33^ stably expressing Venus-mAID-LMNA. Stable single integrants were selected with 0.5 µg/ml puromycin (NLS-TIR1_P2A_NES-TIR1) and 200 µg/ml hygromycin (Venus-mAID-LMNA). For transient transfection with pCS2+_Flag-myc-NES-TIR1 and pIREShygro-Venus-mAID-LMNA, plasmids were co-electroporated into HeLa Kyoto cells according to the manufacturer’s instructions and seeded into 10 cm cell culture dishes. After 24-36 hours, cells were split into 96-well plastic bottom plates (µclear, Greiner Bio-One) and 2.5 mM thymidine (Sigma-Aldrich) was added to pre-synchronize cells at the border of G1 to S-phase. After 24 hours of thymidine arrest, cells were washed and released into fresh media containing 332 nM nocodazole (Sigma-Aldrich) to ensure that cells remain in mitosis due to mitotic checkpoint activation during Venus-mAID-LMNA degradation. Stable HeLa FRT/TO NLS-TIR1_P2A_NES-TIR1/Venus-mAID-LMNA cells were pre-synchronized by the same protocol.

### Plasmid Construction

To generate pAZ132-coPH-oma-1(219-378), pGF06^2^ was amplified with primers oma-1(378) Stop attL2 F + oma-1(219) coPH R and a C-terminal fragment of *oma-1* genomic DNA (aa219-378) was amplified with primers coPH oma-1(219) F + attL2 Stop oma-1(378) R. The PCR fragments were assembled using Gibson Assembly Mix (NEB) and pDonr221-coPH-oma-1(219-378) was then recombined into pAZ132-Gtwy (gift of Barth Grant) using Gateway cloning (Invitrogen).

To generate ss-pCFJ1954-KDEL, the signal sequence of *sel-1* was codon-optimized^53^ and cloned into pCFJ1954 using around-the-world PCR with primers sel-1 ss flex F + sel-1 ss eft-3p R followed by a KLD reaction (NEB). A codon-optimized ZF1 domain from *pie-1* and the KDEL sequence were then cloned into ss-pCFJ1954 using around-the-world PCR with primers coZF1 KDEL Stop attB2 F + coZF1 flex R followed by a KLD reaction.

To generate pIREShygro-Venus-mAID-LMNA, the LMNA open reading frame was amplified from a plasmid encoding GFP-Lamin A^54^ and cloned onto the C-terminus of 3xHA-Venus-mAID^33^ within a pIRES-hygro3 backbone (Clontech).

To localize TIR1 to the cytoplasm, pCS2+_Flag-myc-NES-TIR1 was used (Addgene plasmid #117717)^33^. To localize TIR1 to both the nucleus and cytoplasm, pCAGGs-NLS-TIR1_P2A_NES-TIR1 was used (Addgene plasmid #117699)^37^.

### Worm transformation

FT205, WEH251, WEH399, WEH434, and WEH447 were made by biolistic transformation using a Bio-Rad PDS-1000, as described^55^. The DP38 strain was bombarded with MP322 (gift of Michael Glotzer^21^) to generate FT205. WEH251 was generated by co-bombardment of pGF04^20^ and pJN254 (gift of Jeremy Nance^56^) into the HT1593 strain. HT1593 was bombarded with pGF13^20^ to generate WEH399, with pAZ132-coPH-oma-1(219-378) to generate WEH434, or with ss-pCFJ1954-KDEL to generate WEH447.

### RNAi experiments

RNA interference (RNAi) was performed by feeding worms double-stranded RNA (dsRNA)-expressing bacteria from the L1 larval stage through adulthood at 25°C (60-70 hours) according to established protocols^57^, except *tat-5* RNAi was sometimes fed starting from the L3/L4 stage for only 24 hours. The following RNAi clones were used from available libraries (Source BioScience): *tat-5* (JA:F36H2.1) and *unc-59* (mv_W09C5.2). The *zif-1* F2 RNAi clone was previously described^35^. The RNAi clone for *tsg-101* was designed according to the Cenix clone 49-f7^58^ and dsRNA was transcribed, as described^35^. 1 mg/ml *tsg-101* dsRNA was injected into the gonad of young adult worms 20-26h before analysis. Efficiency of *tsg-101* RNAi was judged by a mild delay in internalization of the AB midbody remnant.

### Light Microscopy

Worm embryos were dissected from gravid adults and mounted in M9 buffer on an agarose pad on a glass slide. For imaging FT205 and FT368, Z-stacks were acquired on a Zeiss AxioImager, 40X 1.3 NA objective, an Axiocam MRM camera, and AxioVision software, as described previously^21^. For N2, FT205, FT368, and XA3502, Z-stacks were acquired for GFP and DIC every minute at room temperature using a Leica DM5500 wide-field fluorescence microscope with a HC PL APO 40X 1.3 NA oil objective lens supplemented with a Leica DFC365 FX CCD camera controlled by LAS AF software. For BV113, WEH02, WEH51, WEH132, WEH142, and WEH248, Z-stacks were acquired sequentially for GFP and mCherry every 20 seconds, as described^20,35^. For WEH251, Z-stacks were acquired for mCherry every 20 seconds or every minute. For strains WEH260 and WEH296, Z-stacks were acquired for mCherry every minute, as described^20^. For WEH399, Z-stacks were acquired for mKate2 and DIC every 30 seconds. For WEH434, Z-stacks were captured for mCherry every 40 seconds. Embryos that arrested during imaging were excluded from analysis. Time-lapse series were analysed using Imaris (Bitplane).

For mammalian cell imaging, a modified DMEM containing 10% (v/v) FBS, 1% (v/v) penicillin-streptomycin, 1% (v/v) Glutamax, and 0.5 µg/ml Amphotericin B without phenol red or riboflavin was used to reduce autofluorescence^59^. DNA was labelled by SiR-Hoechst (Spirochrome) at a final concentration of 50 nM 1-2 hours prior to imaging. Cells were imaged for 6 hours every 5 minutes by automated time-lapse microscopy on an ImageXpress Micro XLS widefield screening microscope (Molecular Devices) equipped with laser-based autofocus, 10 × /0.5 numerical aperture (NA) and 20 × /0.7 NA air objectives (Nikon), a Spectra X light engine (Lumencor), a sCMOS (Andor) camera (binning = 1 or 2), filters for YFP, Texas Red and Cy5 and a stage incubator at 37 °C in a humidified atmosphere of 5% CO_2_. Auxin-dependent degradation was initiated at the beginning of imaging by adding 1-naphthaleneacetic acid sodium salt (NAA, ChemCruz) to a final concentration of 0.5 µM. Cells that died during imaging were excluded from analysis.

### Progeny counts

Single L4 worms from wild type N2 and degron reporter strains were singled onto three 12-well plates. The wells were photographed 3.75 days later on a Leica M80 with a Leica MC120 HD camera. Juvenile L2-L4 and adult progeny were counted using the Cell Counter in Fiji (NIH). For comparison of LMN-1 reporter strains, three L4 XA3502 worms or WEH251 with or without *zif-1* RNAi treatment were put on one 60 mm plate and progeny were counted 3.75 days later. The number of worms with motility deficits (Unc) was reported as a percentage of the total worms counted on each plate. Wells where the mother died (n=10) or the worms clumped (n=12) were excluded from progeny counts.

### Cell cycle timing

To compare the speed of development between control and ZF1-tagged strains, the time from the 3- to the 6-, 12-, and 24-cell stages (one to three subsequent cell cycles in the AB lineage) were calculated from time-lapse series. The time of the AB cell division (3-cell stage) was compared to the timing of the AB daughter (ABx, 6-cell), AB granddaughter (ABxx, 12-cell) and AB great-granddaughter (ABxxx, 24-cell) cell divisions. The cell cycle time was measured from cytokinetic furrow ingression in the AB cell to the ingression of any furrow in the two ABx, four ABxx or eight ABxxx cells. One embryo from FT205 strain was excluded because of cytokinesis defect.

### Tracking

H2B-labeled nuclei were tracked over time using the surface function of Imaris with thresholding to segment objects.

### Quantification of corpse membrane breakdown

Corpse membrane topology was measured using a line scan across the 2PB phagosome in the WEH434 strain. A line with 3-pixel thickness was drawn through the middle of the phagosome using Fiji (NIH) and the mean profile intensity was measured.

### Quantification of nuclei size

The widest cross-section was used to calculate the nuclear area prior to AB cell division. The edge of the P1 nuclear lamina was traced using Fiji (NIH) from fluorescent images of XA3502 and WEH251, as well as WEH251 fed with *zif-1* RNAi.

### Fluorescence intensity measurements

Mean fluorescence intensity of the mCh::PH::CTPD reporter (Fig. 3) was measured in a circle with an area of 0.5 µm^2^ using ImageJ (NIH). Fluorescence intensity of the plasma membrane at the anterior side of the embryo near the first polar body was measured. Fluorescence intensity of the first polar body was measured as an internal control to correct for bleaching. Data are reported as the ratio of the fluorescence intensity of the plasma membrane to that of the polar body.

Mean fluorescence intensity of the ZF1::mCh::CHC-1 reporter (Fig. 4) was measured from images of 4- and 6-cell embryos. GFP::ZF1::PH fluorescence was used to trace a 5.48 µm long two-pixel width line along the plasma membrane using Fiji (NIH). Cell contacts between the anterior (ABa/ABp) or posterior (EMS/P2) cells were measured on the plasma membrane, in addition to the cytoplasm in both neighbouring cells. Data is reported as the mean intensity of the plasma membrane divided by the mean intensity of the cytoplasm.

Mean fluorescence intensity of the LMN-1 reporter (Fig. 7) was measured in a circle with an area of 0.8 µm^2^ using ImageJ (NIH) in AB and later in ABp and its daughter nuclei as well as P1 and later in EMS nuclei. Fluorescence intensity was measured until the LMN-1 marker in ABp daughter nuclei reappeared in *zif-1* RNAi-treated embryos. Similar time range was used in embryos not treated with RNAi. Fluorescence intensity of the P1 and EMS nuclei was measured as an internal control. Data are reported as the ratio of the fluorescence intensity of the AB lineage nuclei to that of the P1 lineage nuclei. The control ratio was normalized to the *zif-1* RNAi ratio to observe changes due to ZIF-1-mediated degradation. A line was fit to three parts of the curve in Excel to calculate the slopes of the sharp drops during mitosis (from −1 to 2.7 min after the start of cytokinesis in the AB cell and from 11.3 to 13 minutes for ABx division) and the mild fluorescence loss during interphase (4 to 11 minutes).

For analysis of Venus-mAID-LMNA degradation (Fig. 8), all images were background-corrected with Fiji using a rolling ball radius of 50 and 100 pixels for images derived from 10x and 20x objectives, respectively. Subsequently, average Venus-mAID-LMNA fluorescence intensities of single cells were measured manually with Fiji, corrected for the remaining background by measuring image regions without cells and smoothed (4 neighbors each size, 4th polynomal order) with Prism 6 (Graphpad). Smoothed intensities of Venus-mAID-LMNA were normalized to the beginning of the time-lapse (t = 0) to monitor degradation during interphase directly after the addition of NAA or normalized to nuclear envelope breakdown (NEBD) to assess mitotic degradation. As transient transfections with NES-TIR1 and Venus-mAID-LMNA did not result in transfection of all cells with both constructs, only cells that expressed Venus-mAID-LMNA and showed degradation after NEBD were considered as double positive and analysed further (Fig. 8a, b, d, f). Average velocities of Venus-mAID-LMNA degradation were expressed as the percent change in Venus-mAID-LMNA per minute during the first 45 minutes of the time lapse for interphase HeLa FRT/TO cells stably expressing NLS-TIR1_P2A_NES-TIR1 and Venus-mAID-LMNA or during 45 minutes before NEBD (interphase) and 45 minutes after NEBD (mitosis) for HeLa cells transiently expressing NES-TIR1 and Venus-mAID-LMNA.

Mean fluorescence intensity of the NMY-2::GFP::ZF1 reporter (Fig. 9) was measured in a circle with an area of 0.5 µm^2^ using ImageJ (NIH), as described previously^35^. Midbody fluorescence was measured from contractile ring closure until the end of the time lapse series or until the midbody was not distinguishable from the cytoplasm. Fluorescence intensity of the first polar body was measured as an internal control. An exponential decay curve was fit to the polar body data using OriginPro (OriginLab) and used to correct for fluorescence loss due to photobleaching. Embryos were excluded if the P0 and AB midbodies were too close to each other (n=4) or if the polar body data did not fit an exponential decay function (n=1). Embryos who arrested development during time-lapse imaging were excluded from measurements (n=3). Embryos treated with tsg-101 RNAi were excluded if the AB midbody remnant internalized by the 6-cell stage (n=6), as the RNAi was judged ineffective. NMY-2 data are reported as the ratio of the fluorescence intensity of the midbody to the expected value of the polar body after cytoplasmic background subtraction. Timing of the onset of degradation of NMY-2::GFP::ZF1 in the cytoplasm was judged by eye by comparing the relative brightness of ABp and EMS using Imaris.

Mean fluorescence intensity of mCh::SP12 and ss::tagRFP-T::ZF1::KDEL (Fig. 10) were measured in a 5.4 µm^2^ circle in 2- to 15-cell embryos using Fiji (NIH). Measurements were taken at the brightest regions near the nucleus in anterior AB lineage cells and posterior P lineage cells. Data is reported as a ratio of the mean fluorescence of the ER in the anterior AB lineage to the posterior P lineage. Images were excluded when the cell identity could not be determined (n=7).

The mean fluorescence of H2B reporters (Fig. S4b) was measured in a circle with an area of 2.6 µm^2^ in the center of interphase nuclei from 2- to 15-cell stage embryos using Fiji (NIH). In older embryos where ZF1::mCh::H2B was no longer visible in AB nuclei, GFP::H2B was used to find the nucleus. Data is reported as a ratio of the mean fluorescence of the anterior AB lineage nuclei to the posterior P lineage nuclei. Images were excluded when AB or P nuclei had condensed mitotic chromosomes (n=29).

Mean fluorescence intensity of LMN-1 reporters for Fig. S4d was measured using a 3 µm line two pixels in width using Fiji (NIH). The line was drawn at the brightest region of the nuclear membrane from images of 2- to 15-cell stage embryos. The intensity was measured in anterior AB cells and posterior P cells, except when P cells were dividing, in which case P sister cells, including E at the 7-cell stage (n=1) or C at the 14-cell stage (n=1) were measured. In older embryos where ZF1-tagged LMN-1was no longer visible in AB nuclei, YFP::LMN-1 was used to trace the nucleus. Embryos were excluded during NEBD of ABx (n=5) or when the nuclei morphology was extremely malformed (n=4) or when the cell identity could not be determined (n=1).

Mean fluorescence intensity of NMY-2 reporters (Fig. S4f) was measured in a circle with an area of 1 µm^2^ in midbodies after abscission in 2- to 15-cell embryos using Fiji (NIH). Data is presented as a ratio of the fluorescence intensity of anterior midbodies to posterior midbodies. In older embryos where ZF1-tagged NMY-2 was no longer visible in AB nuclei, NMY-2::mCh was used to locate the midbody. Images were excluded when midbodies were out of focus (n=3) or endocytosed (n=20) or when the cell identity could not be determined (n=1).

### Fluorescence recovery after photobleaching

Fluorescence recovery after photobleaching (FRAP) was conducted using a Leica SP8 confocal with a HCX PL APO CS 40.0×1.25-NA oil objective with PMT detectors. Half of the ABp nucleus was bleached using the 560 nm laser line. Images were acquired before and after bleaching with intervals of 2 seconds (5 times), 5 seconds (15 times), and 10 seconds (10 times). Fluorescence intensity was measured in three areas of 0.5 µm^2^ on the nuclear envelope within the bleached region and three areas within the nonbleached region using ImageJ (NIH). Data are reported as the ratio of the average intensity in bleached regions to the average intensity in nonbleached regions. Embryos were excluded when focus was lost very early during imaging (n=3). In 2 movies focus was lost during the last two minutes. Those time points were not included in measurements.

### Image processing

For clarity, images were rotated, colorized and the intensity was adjusted using Adobe Photoshop. All images show a single optical section (Z), except for Fig. 5, Fig. 7, Fig. 10, Fig. S2, and Fig. S5. In Fig. 5, six Zs with 1.2 μm steps were maximum projected using Leica LAS X software. In Fig. 7, three Zs with 1.2 μm steps were maximum projected using Leica LAS X. In Fig. 10, two Zs with 1.2 μm steps were maximum projected using Fiji (NIH). In Fig. S2, images where maximum projected to span a region of 2.5 μm using Fiji (NIH). In Fig. S4a, four Zs with 1.2 μm steps were maximum projected using Fiji (NIH). In Fig. S4c, two Zs with 1.2 μm steps were maximum projected using Fiji (NIH). In Fig. S4e, six Zs with 1.2 μm steps were maximum projected using Fiji (NIH). Fig. 5c, 5f, and videos were rotated, colorized, and the intensity was adjusted using Imaris. Several Zs were maximum projected in Imaris for videos. For the left video in video 4, the brightness was adjusted in each frame using Photoshop to compensate for photobleaching.

### Statistical evaluation

Student’s one-tailed *t*-test was used to test statistical significance, except Fisher’s exact test and a chi-squared test were used to test statistical significance of categorical data. A Bonferroni correction was used to adjust for multiple comparisons. Prism 6.0 (Graphpad) was used for statistics and to create graphs for Fig. 8. Venus-mAID-LMNA degradation curves represent single cell data of the indicated number of cells. Differences in average Venus-mAID-LMNA degradation velocities were analysed for significance using an unpaired multi-comparison Kruskal-Wallis test with Dunn’s statistical hypothesis testing presenting multiplicity-adjusted P values. All experiments are representative of at least three independent repeats unless otherwise stated. No randomization or blinding was used in this study.

## Data Availability

The source data underlying Figs 3c, 4e, 7g-h, 8f, 8g, 8h, 9g, 10c and Supplementary Figs 1a-f, 3, 4b, 4d, 4f are provided as a Source Data file. Full data sets generated during and/or analysed during the current study are available from the corresponding author on request.

## Supplemental Figure Legends

**Fig. S1:**
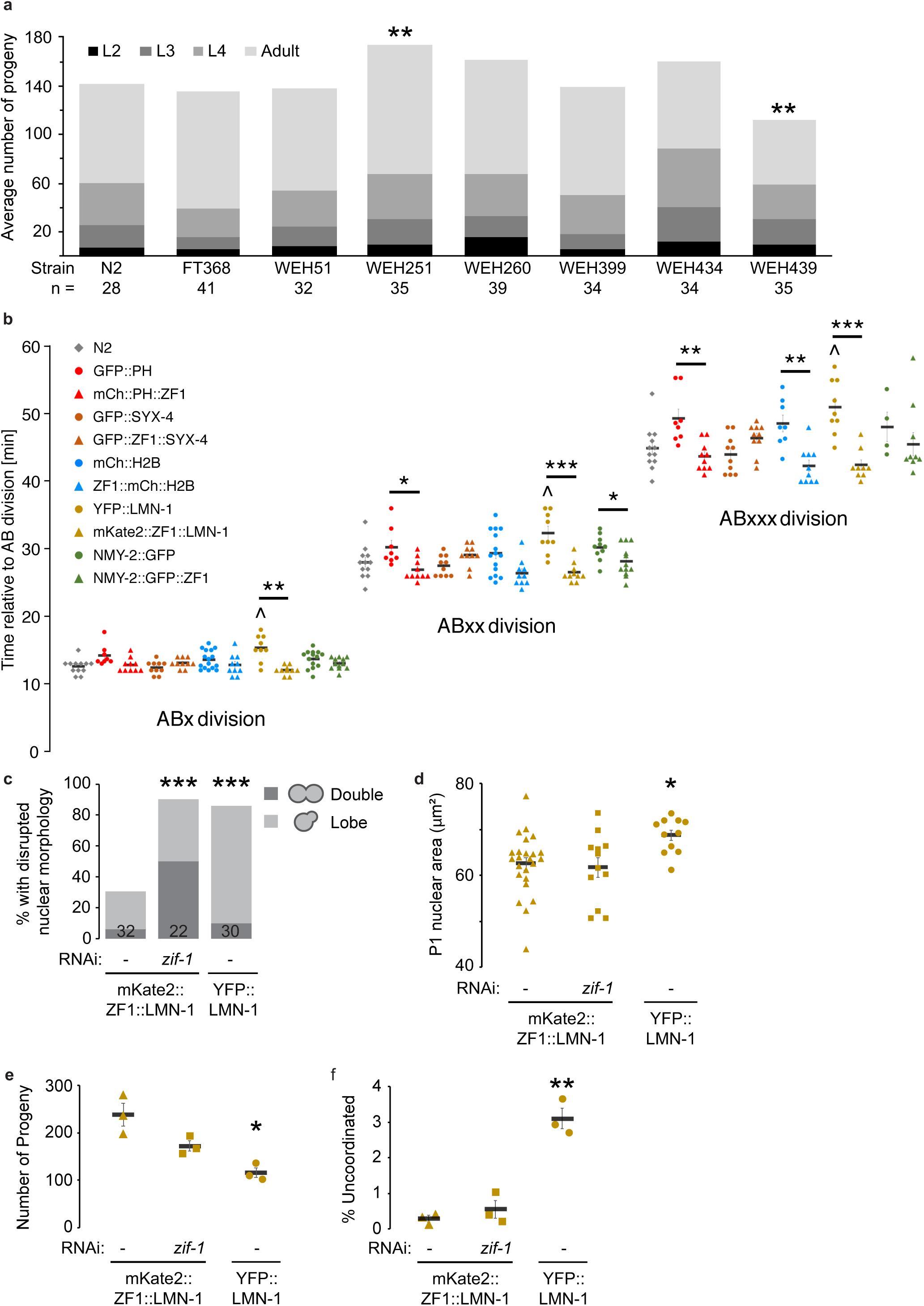
Degron-tagged reporters do not generally disrupt development. a) None of the indicated degron-reporter strains resulted in delays in long-term development (p>0.1, Chi-squared test). WEH251 (mKate2::ZF1::LMN-1) had a significant increase in the total number of progeny in comparison to wild type worms (N2), while WEH439 expressing two degron-tagged reporters (ZF1::mCh::CHC-1; GFP::ZF1::PH) showed a significant reduction in total progeny (**p<0.01, Student’s t-test with Bonferroni correction). The number of wells per worm strain is indicated. b) The timing of subsequent divisions of the anterior AB cell were compared between wild type (grey diamonds, n=12), control reporter strains (circles) and ZF1-tagged reporter strains (triangles), where the same protein was fluorescently-tagged with or without the ZF1 degron. Developmental timing was not significantly delayed after ZF1-tagging the PH domain of PLC1∂1 (red), the syntaxin SYX-4 (brown), a histone H2B (blue), nuclear Lamin LMN-1 (yellow), or non-muscle myosin NMY-2 (green) (p>0.1 using Student’s t-test with Bonferroni correction). Developmental timing was delayed in a strain expressing YFP-tagged LMN-1 without a degron (^p<0.05). Division timing was slower in control fluorescent-tagged strains than their ZF1-tagged counterparts (PH strains: n=10 WEH260 vs. n=8 WEH02 embryos, SYX-4 strains: n=10 FT368 vs. n=10 FT205, H2B strains: n=10 WEH296 vs. n=5 WEH142 or n=3-11 WEH248, LMN-1 strains: n=9-10 WEH251 vs. n=9 XA3502, and NMY-2 strains: n=10-12 WEH51 vs. n=4-14 BV113) Bars represent mean ± SEM. *p<0.05, **p<0.01, ***p<0.001. c) Nuclear morphology was disrupted after LMN-1 overexpression. The penetrance of double nuclei or lobed nuclei was increased after disrupting degron-mediated degradation of mKate2::ZF1::LMN-1 by *zif-1* RNAi (***p<0.001 using Fisher’s exact test). The number of embryos examined are indicated. d) The P1 nucleus was significantly enlarged in the YFP::LMN-1 strain XA3502 (n=11) compared to the degron-tagged strain with (n=12) or without *zif-1* RNAi (n=24). e) The total number of progeny was significantly decreased in the YFP::LMN-1 strain XA3502 compared to the degron-tagged strain with or without *zif-1* RNAi. f) The percentage of uncoordinated (Unc) worms was significantly increased in the YFP::LMN-1 strain XA3502 compared to the degron-tagged strain with or without *zif-1* RNAi. In d-f, bars represent mean ± SEM (*p<0.05, **p<0.01 using Student’s t-test with Bonferroni correction).

**Fig. S2:**
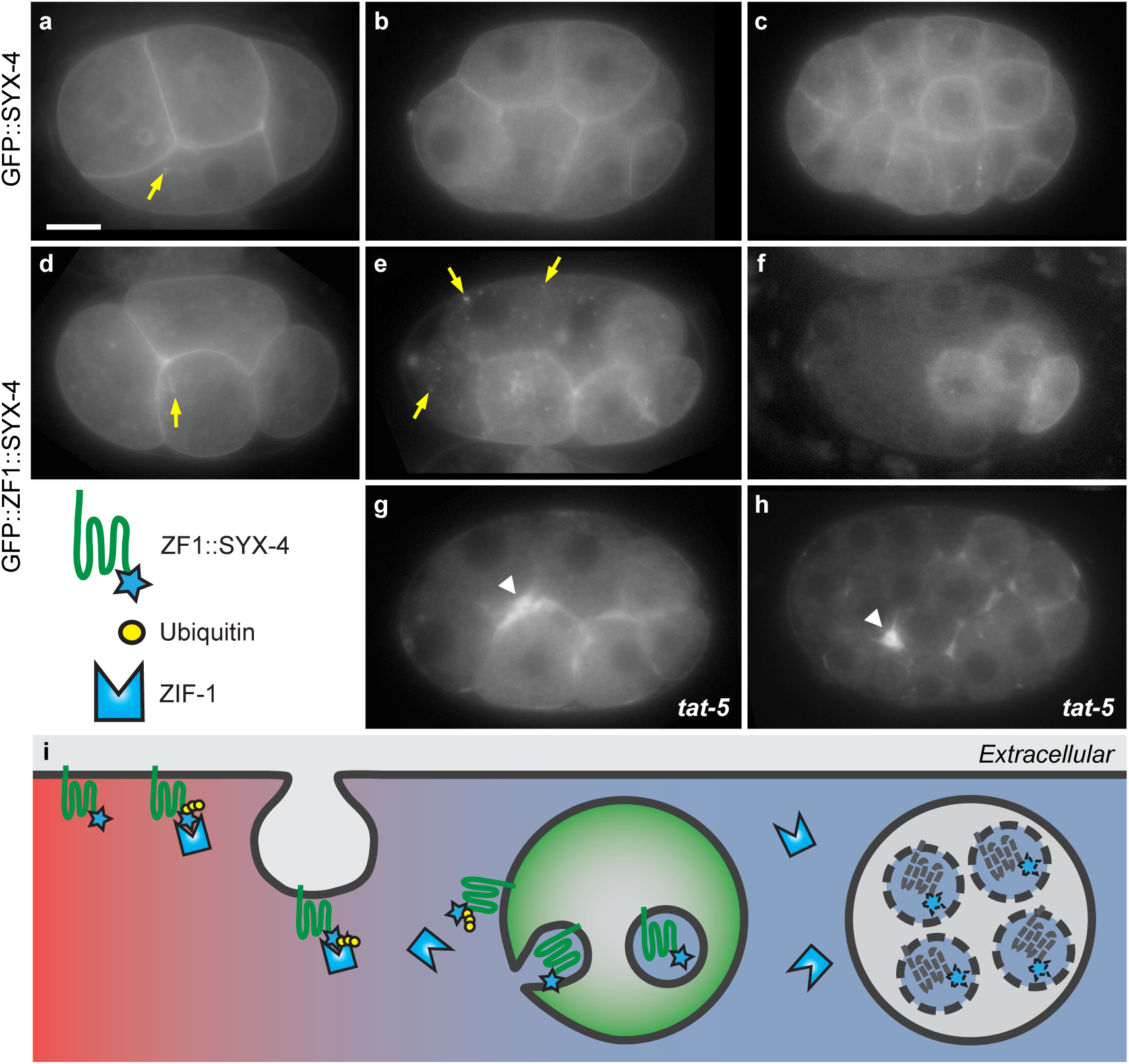
Degron tags drive endocytosis and lysosomal degradation of a transmembrane reporter. a-c) A transmembrane syntaxin localizes to the plasma membrane and endosomes (arrow) in 4-, 8-, and 26-cell GFP::SYX-4-expressing embryos (n=34). Scale bar: 10 µm. d) Prior to ZIF-1 expression, the degron-tagged GFP::ZF1::SYX-4 reporter localizes to the plasma membrane and endosomes (arrow). e) After ZIF-1 expression starts in anterior blastomere (AB) daughter cells, GFP::ZF1::SYX-4 is endocytosed (arrows) and is no longer found in the plasma membrane. f) After endocytosis, GFP::ZF1::SYX-4 vesicles disappear from the anterior cells by the 26-cell stage, consistent with lysosomal degradation (n=43). Anterior is left, dorsal is up. g-h) Embryos treated with *tat-5* RNAi show increased GFP::ZF1::SYX-4 membrane labelling due to accumulated microvesicles (arrowhead) at the 8- and 26-cell stage (n=29). i) A transmembrane protein tagged with the ZF1 degron is recognized by the ubiquitin ligase adaptor ZIF-1 at the plasma membrane and ubiquitinated. Polyubiquitination leads to endocytosis and microautophagy, which results in degradation of reporters on intraluminal vesicles within lysosomes (dotted lines) of degron-tagged fluorescent reporters.

**Fig. S3:**
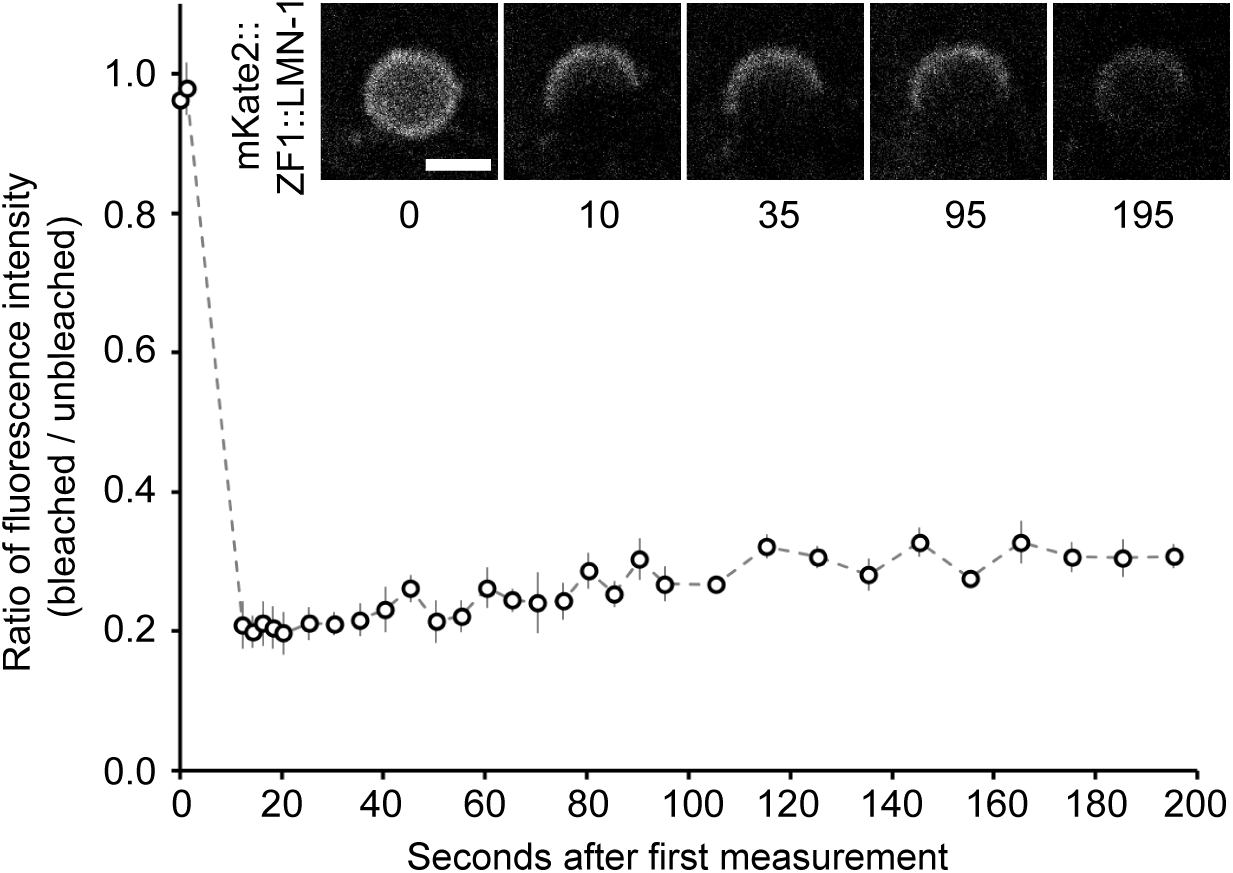
LMN-1 is immobile during degron-mediated degradation. Half of the ABp nuclear lamina expressing mKate2::ZF1::LMN-1was photobleached during the 4-cell stage and imaged. The fluorescence of mKate2::ZF1::LMN-1 does not significantly recover after photobleaching (p>0.3 using Student’s t-test with Bonferroni correction), indicating that its mobility is not increased after ubiquitination (n=3-5). Data are represented as mean ± SEM. Scale bar: 5 µm.

**Fig. S4:**
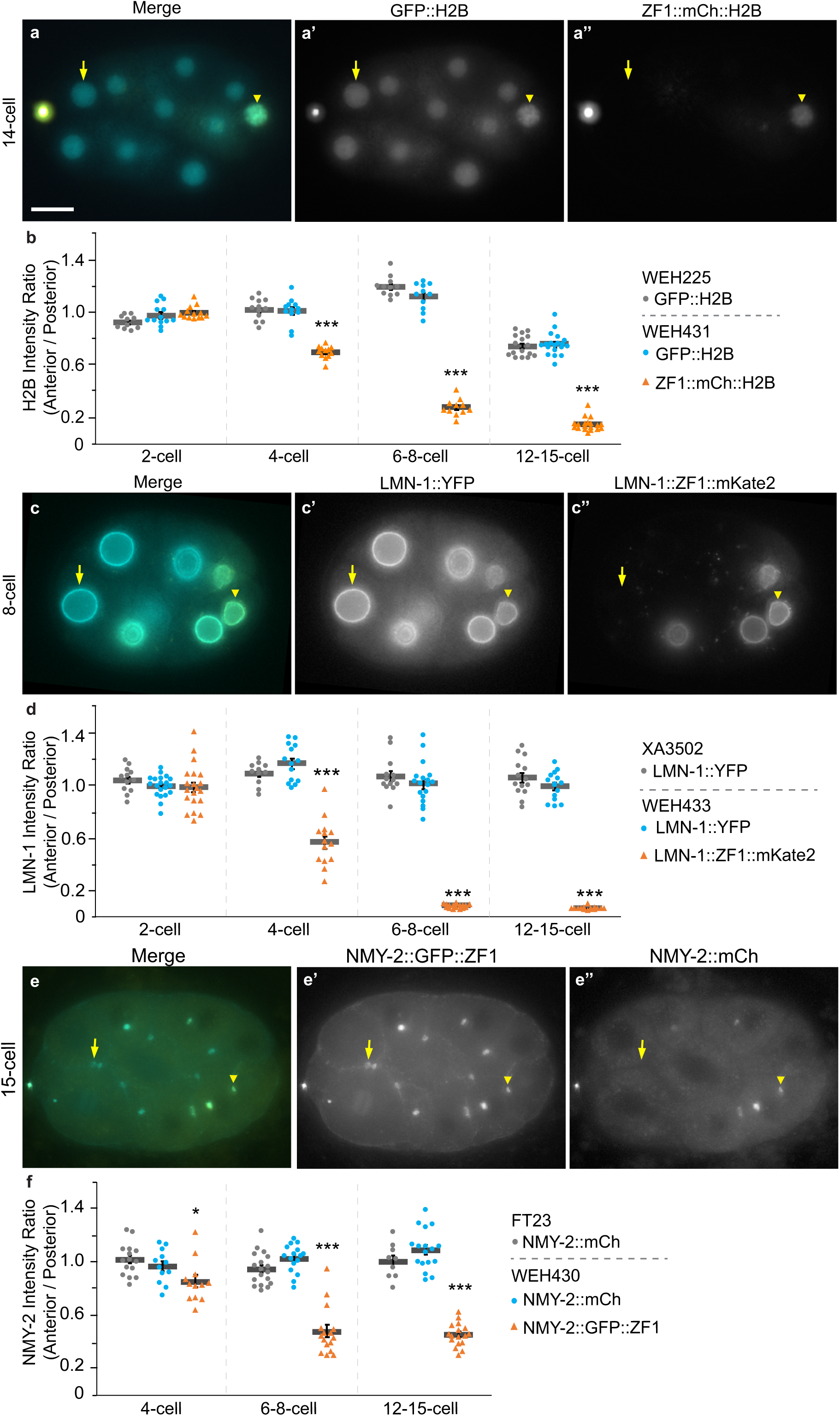
Degron-tagged reporters do not cause degradation of untagged binding partners. a) The WEH431 strain expresses GFP-tagged H2B and ZF1 degron::mCherry-tagged H2B. Scale bar: 10 µm. b) The ratio of the fluorescence between anterior (somatic, arrow) and posterior (germ line, arrowhead) nuclei of the WEH431 strain (n=11-19) was compared with the ratio of the fluorescence of the WEH225 strain (n=10-16) expressing only GFP-tagged H2B at the same cell stage. ZF1::mCh::H2B drops significantly in anterior somatic cells starting during the 4-cell stage, but there is no significant change to GFP::H2B fluorescence in the presence of the degron-tagged H2B reporter (p>0.1). c) The WEH433 strain expresses YFP-tagged LMN-1 and mKate2::ZF1 degron-tagged LMN-1. d) The ratio of the fluorescence between anterior (somatic, arrow) and posterior (germ line, arrowhead) nuclear lamina of the WEH433 strain (n=14-20) was compared with the ratio of the fluorescence of the XA3502 strain (n=11-14) expressing only YFP-tagged LMN-1 at the same cell stage from two independent experiments. mKate2::ZF1::LMN-1 drops significantly in anterior somatic cells from the 4-cell stage, but there is no significant change to YFP::LMN-1 in the presence of the degron-tagged LMN-1 reporter (p>0.3). e) The WEH430 strain expresses mCherry-tagged NMY-2 and GFP::ZF1 degron-tagged NMY-2. f) The ratio of the fluorescence between anterior (somatic, arrow) and posterior (germ line, arrowhead) midbodies of the WEH430 strain (n=12-17) was compared with the ratio of the fluorescence of the FT23 strain (n=9-18) expressing only mCherry-tagged NMY-2 at the same cell stage. NMY-2::GFP::ZF1 drops significantly in anterior somatic cells starting during the 4-cell stage, but there is no significant change in NMY-2::mCherry in the presence of the degron-tagged NMY-2 reporter (p>0.05). Bars represent mean ± SEM (*p<0.05, ***p<0.001 using Student’s t-test with Bonferroni correction).

## Video Legends

Video 1. ZF1 degron reporters facilitate detection of released extracellular vesicles and polar bodies

Time-lapse movies of WEH260 embryos expressing mCh::PH::ZF1 on membranes (cyan) with or without *tat-5* RNAi treatment to induce extracellular vesicle accumulation. In an untreated control embryo, mCh::PH::ZF1 localizes to the plasma membrane, but gradually disappears in somatic cells, starting with the anterior cells. In a *tat-5* RNAi-treated embryo, mCh::PH::ZF1 persists on released extracellular vesicles that accumulate between cells. mCh::PH::ZF1 persists in posterior germ cells, as well as in polar bodies in both control and *tat-5* RNAi-treated embryos. Anterior is to the left, dorsal is up. Time-lapse data were collected every minute for 90 minutes (6 fps). 9 Zs are projected with an interval of 1.2 µm.

Video 2. CTPD degron reporters facilitate detection of released extracellular vesicles and polar bodies

Time-lapse movies of WEH434 embryos expressing mCh::PH::CTPD on membranes (cyan) with or without *tat-5* RNAi treatment to induce extracellular vesicle accumulation. In an untreated control embryo, mCh::PH::CTPD localizes to the plasma membrane, but gradually disappears during the first cell division. In a *tat-5* RNAi-treated embryo, mCh::PH::CTPD persists on released extracellular vesicles that accumulate between cells. mCh::PH::CTPD persists in polar bodies in both control and *tat-5* RNAi-treated embryos. Anterior is to the left, dorsal is up. Time-lapse data were collected every 40 seconds for 27 minutes (6 fps). 12 Zs are projected for the control embryo with an interval of 1 µm, while 10 Zs are projected for the *tat-5(RNAi)* embryo with an interval of 1.2 µm.

Video 3. Degron reporters enable tracking of a phagocytosed cell corpse

Time-lapse movies of WEH142 embryos expressing mCh::H2B and WEH339 embryos expressing ZF1::mCh::H2B in nuclei, including both polar bodies (arrowheads). It is harder to follow the corpse of the second polar body labelled with mCh::H2B, due to the fluorescence of nearby nuclei. ZF1::mCh::H2B is degraded in sequential sets of somatic nuclei, facilitating tracking of the polar bodies as they remain the only fluorescent structures in the anterior half of the embryo. Anterior is to the left, dorsal is up, and the embryos are within the dotted oval. Time-lapse data were collected every 20 seconds for WEH142 and every 30 seconds for WEH339 for 40 minutes in total (playing at 15 fps and 10 fps, respectively). 11 Zs with 1.2 µm intervals are projected.

Video 4. Degron reporters reveal nuclear envelope dynamics during cell division

Time-lapse movies of WEH251 embryos expressing mKate2::ZF1::LMN-1 to mark the nuclear lamina. In control embryos, the reporter gradually loses fluorescence intensity in anterior somatic cells during interphase, followed by a sharp drop of fluorescence in dividing cells when nuclear envelope breakdown occurs. When the ubiquitin ligase adaptor ZIF-1 is knocked down, the mKate2::ZF1::LMN-1 fluorescence is stable in all cells. Anterior is to the left, dorsal is up. Time-lapse data were collected every 20 seconds for 21-22 minutes (6 fps). 8 Zs with 1 µm intervals are projected.

Video 5. LMNA is protected from NES-TIR1 during interphase

Representative time-lapse series of a HeLa cell transiently expressing NES-TIR1 and Venus-mAID-LMNA. 0.5 mM NAA was added at t = 0 to induce auxin-dependent degradation. During interphase, Venus-mAID-LMNA in the nuclear lamina is protected from ubiquitination by cytosolic NES-TIR1 and fluorescence persists in the presence of NAA. Data were collected every 5 minutes. Scale bar: 30 µm.

Video 6. LMNA is accessible to NES-TIR1 during mitosis

Representative time-lapse series of a HeLa cell transiently expressing NES-TIR1 and Venus-mAID-LMNA entering mitosis (same cell as Video 5). After nuclear envelope breakdown (NEBD, t = 0), Venus-mAID-LMNA is accessible by cytosolic NES-TIR1 and fluorescence rapidly disappears. Data were collected every 5 minutes. Scale bar: 30 µm.

Video 7. LMNA is accessible to NLS-TIR1 during interphase

Interphase HeLa cell stably expressing NLS-TIR1 and NES-TIR1 from a single mRNA in addition to Venus-mAID-LMNA. After treatment with 0.5 mM NAA (t = 0), Venus-mAID-LMNA fluorescence rapidly disappears during interphase. Data were collected every 5 minutes. Scale bar: 30 µm.

## Supplemental Tables

**Table S1:**
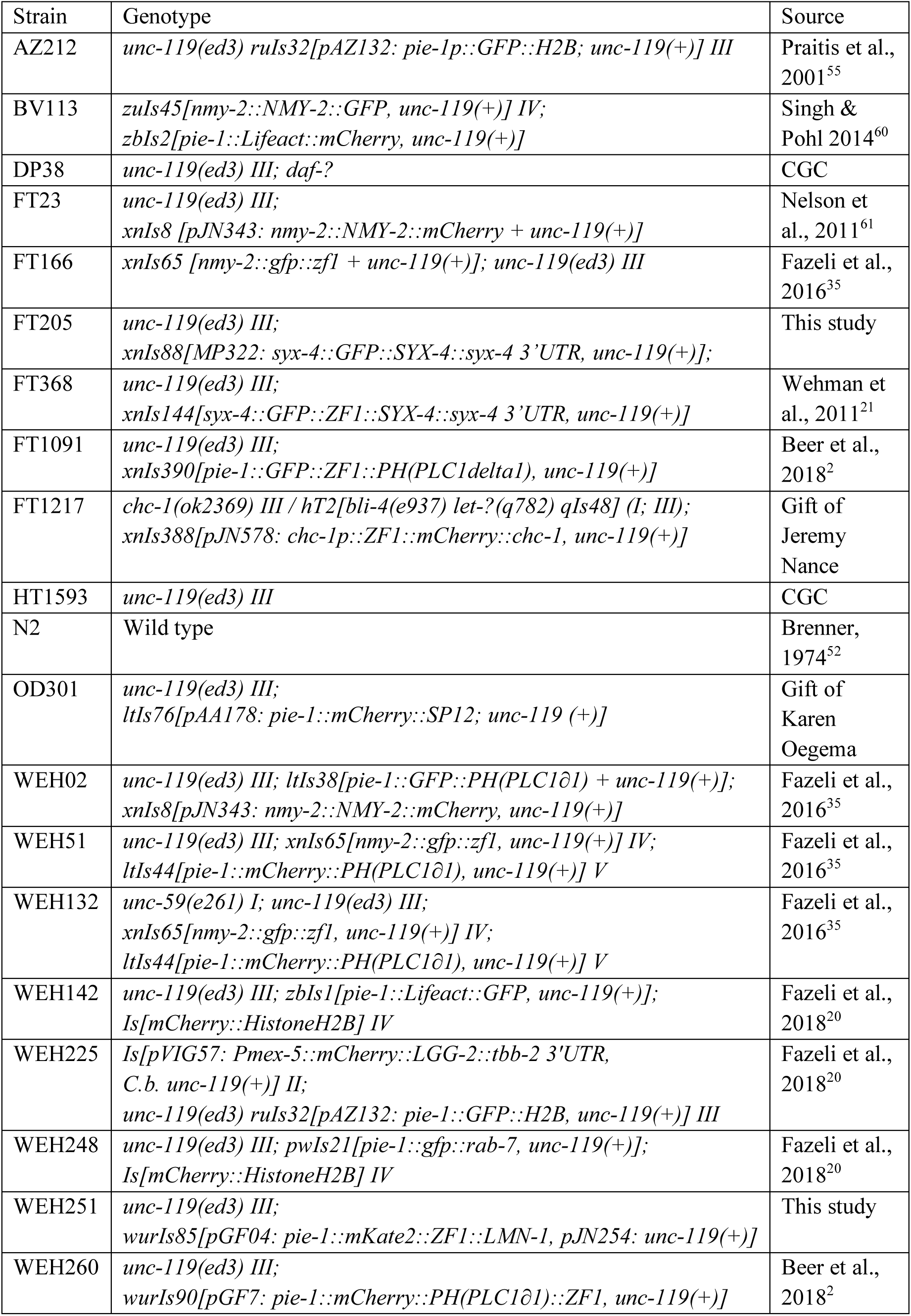
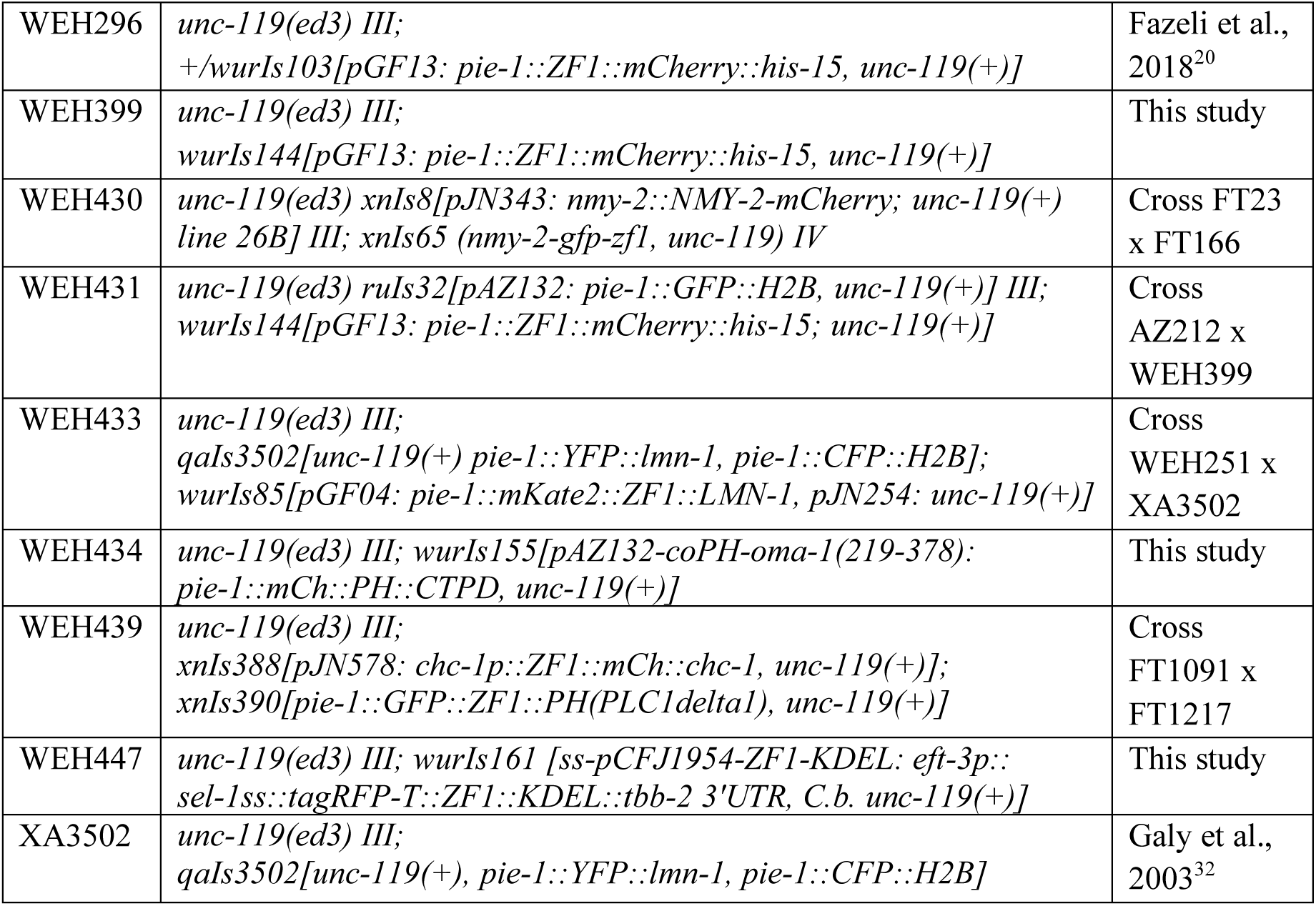
Strains used in this study.

**Table S2:**
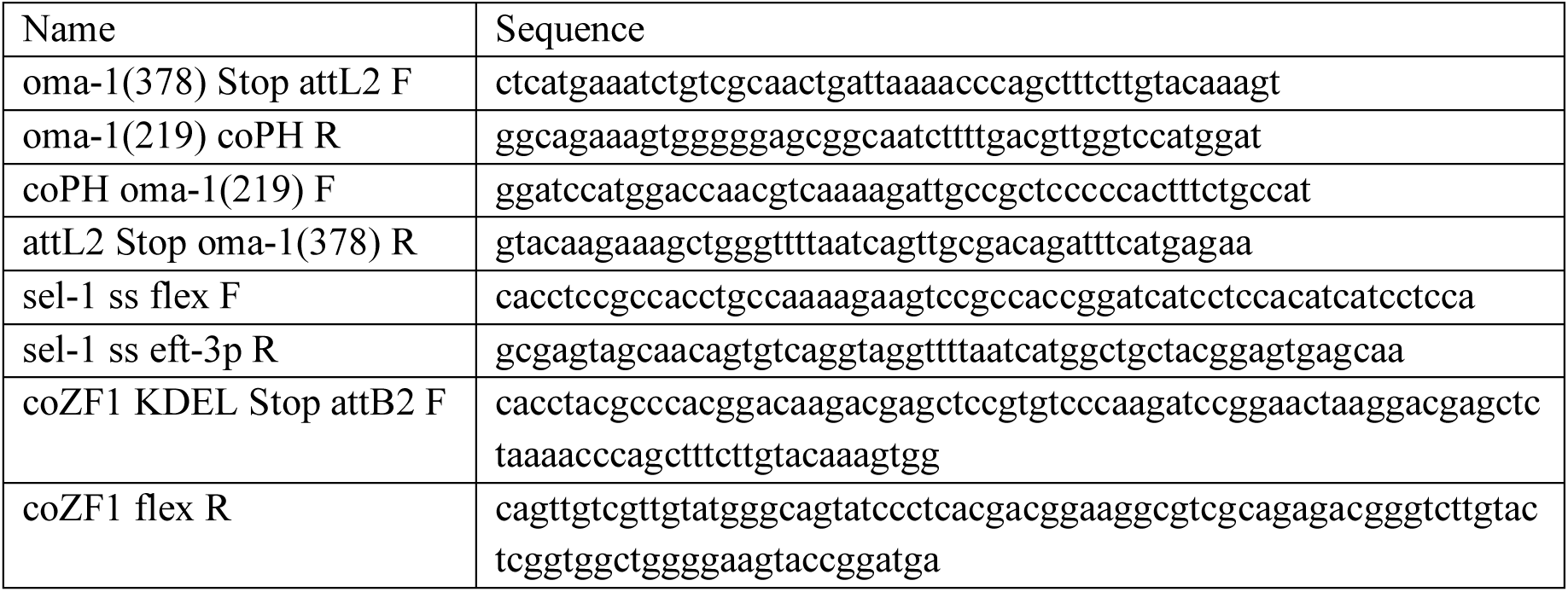
Primers used in this study.

## References

1. Edidin, M. Lipids on the frontier: A century of cell-membrane bilayers. Nat. Rev. Mol. Cell Biol. 4, 414–418 (2003).

2. Beer, K. B. et al. Extracellular vesicle budding is inhibited by redundant regulators of TAT-5 flippase localization and phospholipid asymmetry. Proc. Natl. Acad. Sci. 115, E1127–E1136 (2018).

3. Renard, H.-F., Johannes, L. & Morsomme, P. Increasing Diversity of Biological Membrane Fission Mechanisms. Trends Cell Biol. 28, 274–286 (2018).

4. Huang, B., Babcock, H. & Zhuang, X. Breaking the diffraction barrier: Super-resolution imaging of cells. Cell 143, 1047–1058 (2010).

5. Várnai, P. & Balla, T. Visualization and manipulation of phosphoinositide dynamics in live cells using engineered protein domains. Pflugers Arch. Eur. J. Physiol. 455, 69–82 (2007).

6. Flannagan, R. S., Jaumouillé, V. & Grinstein, S. The Cell Biology of Phagocytosis. Annu. Rev. Pathol. Mech. Dis. 7, 61–98 (2012).

7. Schermelleh, L. et al. Super-resolution microscopy demystified. Nat. Cell Biol. 21, 72–84 (2019).

8. Beer, K. B. & Wehman, A. M. Mechanisms and functions of extracellular vesicle release in vivo—What we can learn from flies and worms. Cell Adhes. Migr. 11, 135–150 (2017).

9. Van Niel, G., D’Angelo, G. & Raposo, G. Shedding light on the cell biology of extracellular vesicles. Nat. Rev. Mol. Cell Biol. 19, 213–228 (2018).

10. Lorenz, H., Hailey, D. W. & Lippincott-Schwartz, J. Fluorescence protease protection of GFP chimeras to reveal protein topology and subcellular localization. Nat. Methods 3, 205 (2006).

11. Natsume, T. & Kanemaki, M. T. Conditional Degrons for Controlling Protein Expression at the Protein Level. Annu. Rev. Genet. 51, 83–102 (2017).

12. Foot, N., Henshall, T. & Kumar, S. Ubiquitination and the Regulation of Membrane Proteins. Physiol. Rev. 97, 253–281 (2017).

13. DeRenzo, C., Reese, K. J. & Seydoux, G. Exclusion of germ plasm proteins from somatic lineages by cullin-dependent degradation. Nature 424, 685–689 (2003).

14. Oldenbroek, M. et al. Multiple RNA-binding proteins function combinatorially to control the soma-restricted expression pattern of the E3 ligase subunit ZIF-1. Dev. Biol. 363, 388–398 (2012).

15. Armenti, S. T., Lohmer, L. L., Sherwood, D. R. & Nance, J. Repurposing an endogenous degradation system for rapid and targeted depletion of C. elegans proteins. Development 141, 4640–4647 (2014).

16. Nishi, Y. & Lin, R. DYRK2 and GSK-3 phosphorylate and promote the timely degradation of OMA-1, a key regulator of the oocyte-to-embryo transition in C. elegans. Dev. Biol. 288, 139–149 (2005).

17. Du, Z., He, F., Yu, Z., Bowerman, B. & Bao, Z. E3 ubiquitin ligases promote progression of differentiation during C. elegans embryogenesis. Dev. Biol. 398, 267–279 (2015).

18. Nishimura, K., Fukagawa, T., Takisawa, H., Kakimoto, T. & Kanemaki, M. An auxin-based degron system for the rapid depletion of proteins in nonplant cells. Nat. Methods 6, 917–922 (2009).

19. Dreher, K. A., Brown, J., Saw, R. E. & Callis, J. The Arabidopsis Aux/IAA protein family has diversified in degradation and auxin responsiveness. Plant Cell 18, 699–714 (2006).

20. Fazeli, G., Stetter, M., Lisack, J. N. & Wehman, A. M. C. elegans Blastomeres Clear the Corpse of the Second Polar Body by LC3-Associated Phagocytosis. Cell Rep. 23, 2070–2082 (2018).

21. Wehman, A. M., Poggioli, C., Schweinsberg, P., Grant, B. D. & Nance, J. The P4-ATPase TAT-5 inhibits the budding of extracellular vesicles in C. elegans embryos. Curr. Biol. 21, 1951–1959 (2011).

22. Jantsch-Plunger, V. & Glotzer, M. Depletion of syntaxins in the early Caenorhabditis elegans embryo reveals a role for membrane fusion events in cytokinesis. Curr. Biol. 9, 738–745 (1999).

23. Beckett, K. et al. Drosophila S2 cells secrete wingless on exosome-like vesicles but the wingless gradient forms independently of exosomes. Traffic 14, 82–96 (2013).

24. Choi, D. S. et al. Proteomic analysis of microvesicles derived from human colorectal cancer cells. J. Proteome Res. 6, 4646–4655 (2007).

25. Benenati, G., Penkov, S., Müller-Reichert, T., Entchev, E. V & Kurzchalia, T. V. Two cytochrome P450s in Caenorhabditis elegans are essential for the organization of eggshell, correct execution of meiosis and the polarization of embryo. Mech. Dev. 126, 382–393 (2009).

26. Florey, O., Kim, S. E., Sandoval, C. P., Haynes, C. M. & Overholtzer, M. Autophagy machinery mediates macroendocytic processing and entotic cell death by targeting single membranes. Nat. Cell Biol. 13, 1335–1343 (2011).

27. Chen, D. et al. Clathrin and AP2 Are Required for Phagocytic Receptor-Mediated Apoptotic Cell Clearance in Caenorhabditis elegans. PLoS Genet. 9, 1–18 (2013).

28. Levin, R., Grinstein, S. & Canton, J. The life cycle of phagosomes: formation, maturation, and resolution. Immunol. Rev. 273, 156–179 (2016).

29. Guven-Ozkan, T., Robertson, S. M., Nishi, Y. & Lin, R. zif-1 translational repression defines a second, mutually exclusive OMA function in germline transcriptional repression. Development 137, 3373–3382 (2010).

30. Gerace, L. & Tapia, O. Messages from the voices within: regulation of signaling by proteins of the nuclear lamina. Curr. Opin. Cell Biol. 52, 14–21 (2018).

31. Ungricht, R. & Kutay, U. Mechanisms and functions of nuclear envelope remodelling. Nat. Rev. Mol. Cell Biol. 18, 229–245 (2017).

32. Galy, V., Askjaer, P., Franz, C., López-Iglesias, C. & Mattaj, I. W. MEL-28, a Novel Nuclear-Envelope and Kinetochore Protein Essential for Zygotic Nuclear-Envelope Assembly in C. elegans. Curr. Biol. 16, 1748–1756 (2006).

33. Daniel, K. et al. Conditional control of fluorescent protein degradation by an auxin-dependent nanobody. Nat. Commun. 9, (2018).

34. Green, R. A. et al. The midbody ring scaffolds the abscission machinery in the absence of midbody microtubules. J. Cell Biol. 203, 505–520 (2013).

35. Fazeli, G., Trinkwalder, M., Irmisch, L. & Wehman, A. M. C. elegans midbodies are released, phagocytosed and undergo LC3-dependent degradation independent of macroautophagy. J. Cell Sci. 129, 3721–3731 (2016).

36. König, J., Frankel, E. B., Audhya, A. & Müller-Reichert, T. Membrane remodeling during embryonic abscission in *Caenorhabditis elegans*. J. Cell Biol. 216, 1277–1286 (2017).

37. Teasdale, R. D. & Jackson, M. R. Signal-Mediated Sorting of Membrane Proteins Between the Endoplasmic Reticulum and the Golgi Apparatus. Annu. Rev. Cell Dev. Biol. 12, 27–54 (2002).

38. Yamaguchi, N., Colak-Champollion, T. & Knaut, H. zGrad is a nanobody-based degron system that inactivates proteins in zebrafish. Elife 8, (2019).

39. Caussinus, E., Kanca, O. & Affolter, M. Fluorescent fusion protein knockout mediated by anti-GFP nanobody. Nat. Struct. & Mol. Biol. 19, 117 (2011).

40. Baudisch, B., Pfort, I., Sorge, E. & Conrad, U. Nanobody-Directed Specific Degradation of Proteins by the 26S-Proteasome in Plants. Front. Plant Sci. 9, 130 (2018).

41. Wang, S. et al. A toolkit for GFP-mediated tissue-specific protein degradation in C. elegans. Development 144, 2694–2701 (2017).

42. Zhang, L., Ward, J. D., Cheng, Z. & Dernburg, A. F. The auxin-inducible degradation (AID) system enables versatile conditional protein depletion in C. elegans. Development 142, 4374–4384 (2015).

43. Blondel, M. Nuclear-specific degradation of Far1 is controlled by the localization of the F-box protein Cdc4. EMBO J. 19, 6085–6097 (2002).

44. Yamada, T., Yang, Y. & Bonni, A. Spatial organization of ubiquitin ligase pathways orchestrates neuronal connectivity. Trends Neurosci. 36, 218–226 (2013).

45. Shin, Y. J. et al. Nanobody-targeted E3-ubiquitin ligase complex degrades nuclear proteins. Sci. Rep. 5, 14269 (2015).

46. Bobrie, A., Colombo, M., Krumeich, S., Raposo, G. & Théry, C. Diverse subpopulations of vesicles secreted by different intracellular mechanisms are present in exosome preparations obtained by differential ultracentrifugation. J. Extracell. Vesicles 1, (2012).

47. Kowal, J. et al. Proteomic comparison defines novel markers to characterize heterogeneous populations of extracellular vesicle subtypes. Proc. Natl. Acad. Sci. 113, E968–E977 (2016).

48. Hell, S. W. Microscopy and its focal switch. Nat. Methods 6, 24–32 (2009).

49. Spiegelhalter, C., Laporte, J. F. & Schwab, Y. Correlative Light and Electron Microscopy: From Live Cell Dynamic to 3D Ultrastructure. in Electron Microscopy: Methods and Protocols (ed. Kuo, J.) 485–501 (Humana Press, 2014). doi:10.1007/978-1-62703-776-1_21

50. Jones, R. D. & Gardner, R. G. Protein quality control in the nucleus. Curr Opin Cell Biol 40, 81–89 (2016).

51. Schrader, E. K., Harstad, K. G. & Matouschek, A. Targeting proteins for degradation. Nat. Chem. Biol. 5, 815–822 (2009).

52. Brenner, S. The genetics of Caenorhabditis elegans. Genetics 77, 71–94 (1974).

53. Redemann, S. et al. Codon adaptation-based control of protein expression in C. elegans. Nat. Methods 8, 250–252 (2011).

54. Broers, J. L. et al. Dynamics of the nuclear lamina as monitored by GFP-tagged A-type lamins. J. Cell Sci. 112 (Pt 2, 3463–75 (1999).

55. Praitis, V., Casey, E., Collar, D. & Austin, J. Creation of low-copy integrated transgenic lines in Caenorhabditis elegans. Genetics 157, 1217–1226 (2001).

56. Nance, J., Munro, E. M. & Priess, J. R. C. elegans PAR-3 and PAR-6 are required for apicobasal asymmetries associated with cell adhesion and gastrulation. Development 130, 5339–5350 (2003).

57. Fraser, A. G. et al. Functional genomic analysis of C. elegans chromosome I by systematic RNA interference. Nature 408, 325–330 (2000).

58. Sönnichsen, B. et al. Full-genome RNAi profiling of early embryogenesis in Caenorhabditis elegans. Nature 434, 462–469 (2005).

59. Schmitz, M. H. A. et al. Live-cell imaging RNAi screen identifies PP2A – B55 α and importin-β 1 as key mitotic exit regulators in human cells. Nat. Cell Biol. 12, 886–93 (2013).

60. Singh, D. & Pohl, C. Coupling of Rotational Cortical Flow, Asymmetric Midbody Positioning, and Spindle Rotation Mediates Dorsoventral Axis Formation in C. elegans. Dev. Cell 28, 253–267 (2014).

61. Nelson, M. D. et al. A bow-tie genetic architecture for morphogenesis suggested by a genome-wide RNAi screen in Caenorhabditis elegans. PLoS Genet. 7, e1002010 (2011).

